# Metabolic resilience rules sex-specific pain recovery during hormonal aging: a multi-omics analysis of neuropathy in mice

**DOI:** 10.1101/2025.06.17.659987

**Authors:** Sara Marinelli, Claudia Rossi, Luisa Pieroni, Giacomo Giacovazzo, Valentina Vacca, Federica De Angelis, Ilaria Cicalini, Valentina Mastrorilli, Chiara Parisi, Zuleyha Nihan Yurtsever, Domenico Ciavardelli, Roberto Coccurello

**Author notes:** Correspondence: Sara Marinelli; Roberto Coccurello. Senior author (Sara Marinelli, Roberto Coccurello).

## Abstract

Biological aging and sex interact to shape systemic metabolism, yet their role in chronic pain resolution remains unexplored. We hypothesized that metabolic resilience — the ability to flexibly switch fuel sources and maintain energy homeostasis — rules successful recovery from nerve injury in a sex-dependent manner during aging.

In 12-month-old male and female mice — corresponding to the perimenopausal phase in females and the onset of hormonal decline in both sexes — we induced sciatic nerve chronic constriction injury and performed multi-omics profiling during Wallerian degeneration, a phase known to trigger long-term neurobiological remodeling. Aging females exhibited early activation of fatty acid oxidation, increased resting energy expenditure, upregulation of mitochondrial redox enzymes and circulating progesterone and corticosterone. Proteomic and metabolomic analysis revealed pentose phosphate pathway enrichment and gluconeogenesis, supporting redox balance and metabolic flexibility. Conversely, males displayed persistent glycolytic reliance, long-chain acylcarnitine accumulation, suppression of adiponectin and PPARγ, indicating metabolic inflexibility.

Longitudinal behavioral analysis revealed that aging females recovered earlier and more fully than aging males, reversing the pattern previously shown in our adult mouse study, where females developed persistent pain and males recovered rapidly. These patterns highlight a non-linear, sex-specific interaction between biological aging and injury response, where hormonal decline reprograms the metabolic trajectory and reshapes pain outcomes.

Metabolic resilience governs sex-specific recovery following nerve injury by directing early systemic adaptations that precede and predict long-term pain trajectories. These results define mechanistically anchored, sex- and age-specific biomarkers, and propose preclinical targets for timely, personalized interventions in age-associated neuropathic pain.

## INTRODUCTION

Peripheral neuropathy (PN) is a highly prevalent condition in older adults, affecting up to 30% of individuals in certain populations [1]. Its incidence increases with age and is disproportionately higher in peri- and postmenopausal women, likely due to the loss of estrogen-mediated neuroprotection and the resulting impairment in neural and immune homeostasis [2,3]. Clinically, PN involves sensory, motor, and autonomic dysfunctions that significantly impair quality of life. Mounting evidence indicates that age-related metabolic and inflammatory changes, coupled with altered body composition—such as increased central adiposity and sarcopenia—contribute to PN development [1,4,5]. In this context, biological sex and aging act as modulators of systemic energy homeostasis, potentially influencing the trajectory of neuropathic pain and its resolution [6].

We and others have previously shown that adipose tissue (AT) is not only an energy storage depot but also a dynamic endocrine organ, modulating both inflammation and nerve regeneration through the secretion of adipokines such as leptin and adiponectin [7,3]. In postmenopausal women, estrogen loss is associated with increased PN prevalence, which has been linked to impaired immune regulation and enhanced oxidative stress [8,9]. Estrogens support Schwann cell (SC) activity and protect peripheral nerves by preserving redox balance and glial function [8,9]. In adult female mice, peripheral nerve injury disrupts metabolic homeostasis, leading to early alterations in lipid metabolism, reduced energy expenditure, and dysregulated steroid secretion from AT, all of which contribute to impaired glycemic and insulin responses [3].

In contrast, male mice display a distinct metabolic response to nerve injury, characterized by enhanced glycolytic flux, lower energy expenditure, and altered fatty acid handling. Notably, AT in males remains functionally responsive, supporting mitochondrial adaptation, oxidative stress compensation, and neuro-regenerative signaling through adiponectin and PPARγ pathways [9–12]. The reduced regenerative activity of SCs in females further delays axonal repair, revealing a complex interplay between sex hormones, glial responses, and metabolic plasticity.

Emerging data also emphasize the importance of AT innervation in maintaining systemic metabolic health. PN involving subcutaneous AT (scAT) disrupts nerve-adipose crosstalk, impairing substrate mobilization and endocrine signaling. Both myelinated and unmyelinated scAT fibers exhibit high plasticity, which is lost in pathological contexts such as neuropathy [13,14]. Dysfunctional SCs in scAT contribute to demyelinating phenotypes, underscoring their dual role in nerve maintenance and injury response [14,15].

Although it is well recognized that aging alters peripheral nerve physiology [4,16,17], the mechanistic drivers of neuropathic pain onset, persistence, and resolution—particularly in the context of age and sex—remain poorly defined. Prior studies have proposed that nerve injury acts as a metabolic switch, triggering energy-intensive repair programs that must be tightly regulated to ensure functional recovery [3,18–20]. In this framework, metabolic inflexibility may serve as an early determinant of pain chronification, particularly in aging organisms where the ability to adapt energy substrates is compromised.

Here, we hypothesize that biological sex modulates the metabolic trajectory of nerve injury response in aging, defining distinct pain outcomes. We further propose that early metabolic rewiring — especially within adipose tissue — governs resilience or vulnerability to chronic pain, offering a window for biomarker-based stratification and sex-specific therapeutic targeting.

To test this, we employed a mouse model of sciatic nerve injury in 12-month-old male and female mice, representing the perimenopausal and early androgen-decline stage, and integrated behavioral, calorimetric, imaging, molecular, and multi-omics analyses. Our approach aimed to define mechanistically anchored, sex-dependent metabolic pathways involved in the early response to nerve injury and to identify predictive markers of long-term pain trajectories. By elucidating these processes, this work supports the development of personalized interventions to prevent or reverse chronic neuropathic pain in aging individuals.

## Methods

### Murine Models of Chronic Constriction Injury-Induced Neuropathic Pain (NeP)

Male and female CD1(ICR)CD1 mice, approximately 12 months old, were obtained from Charles River Labs (Como, Italy) or the European Mouse Mutant Archive – EMMA Infrafrontier (Monterotondo RM, Italy) for this study. The animals were housed under standard conditions in transparent plastic cages, in groups of four, with sawdust bedding, following a 12-hour light/dark cycle (07:00 AM – 07:00 PM). Food and water were provided ad libitum. After behavioral testing, female estrous cycle stages were determined using vaginal smears. Since no significant differences were observed in behavioral responses across different estrous phases, all female mice were included in the same experimental group regardless of hormonal status. All procedures strictly adhered to European and Italian national regulations governing the use of animals in research (Legislative Decree No. 26 of 04/03/2014, implementing European Communities Council Directive 10/63/EU) and were approved by the Italian Ministry of Health (authorization numbers 32/2014PR and 164/2024PR). To induce neuropathic pain, the Chronic Constriction Injury (CCI) model was employed, adapted for mice from the original method described by Bennett and Xie [28] Surgery was performed under anesthesia using a 1:1 mixture of Rompun (Bayer, 20 mg/ml; 0.5 ml/kg) and Zoletil (100 mg/ml; 0.5 ml/kg). A longitudinal skin incision (∼1.5 cm) was made to expose the mid-portion of the right sciatic nerve. Three loosely tied ligatures (7-0 chromic gut, Ethicon, Rome, Italy) were placed around the nerve to induce partial constriction. The incision was then closed using 4-0 silk sutures. From this point forward, the injured right hindpaw will be referred to as the “ipsilateral paw,” while the uninjured left hindpaw will be termed the “contralateral paw.” Mice underwent behavioral assessments (in vivo experiments) or were sacrificed for tissue collection (ex vivo experiments) before CCI surgery to obtain baseline reference values. The experimental timeline is detailed in Figure 1. The study included male and female CD1 mice, evaluated before and at various time points after CCI surgery in in vivo experiments. The first cohort of mice underwent a series of assessments, arranged from the least to the most stressful to minimize experimental interference: allodynia evaluation, glycemia and triglyceride (TG) measurements, body temperature recording, and cold exposure followed by body temperature measurement. The second cohort was designated for indirect calorimetry experiments. For *ex vivo* analyses, each time point represented a separate group of male or female CD1 mice, as animals were sacrificed for tissue collection. The number of animals (N) in each group is specified in the figure legends, corresponding to the respective experimental conditions and statistical analyses.

**Figure 1.**
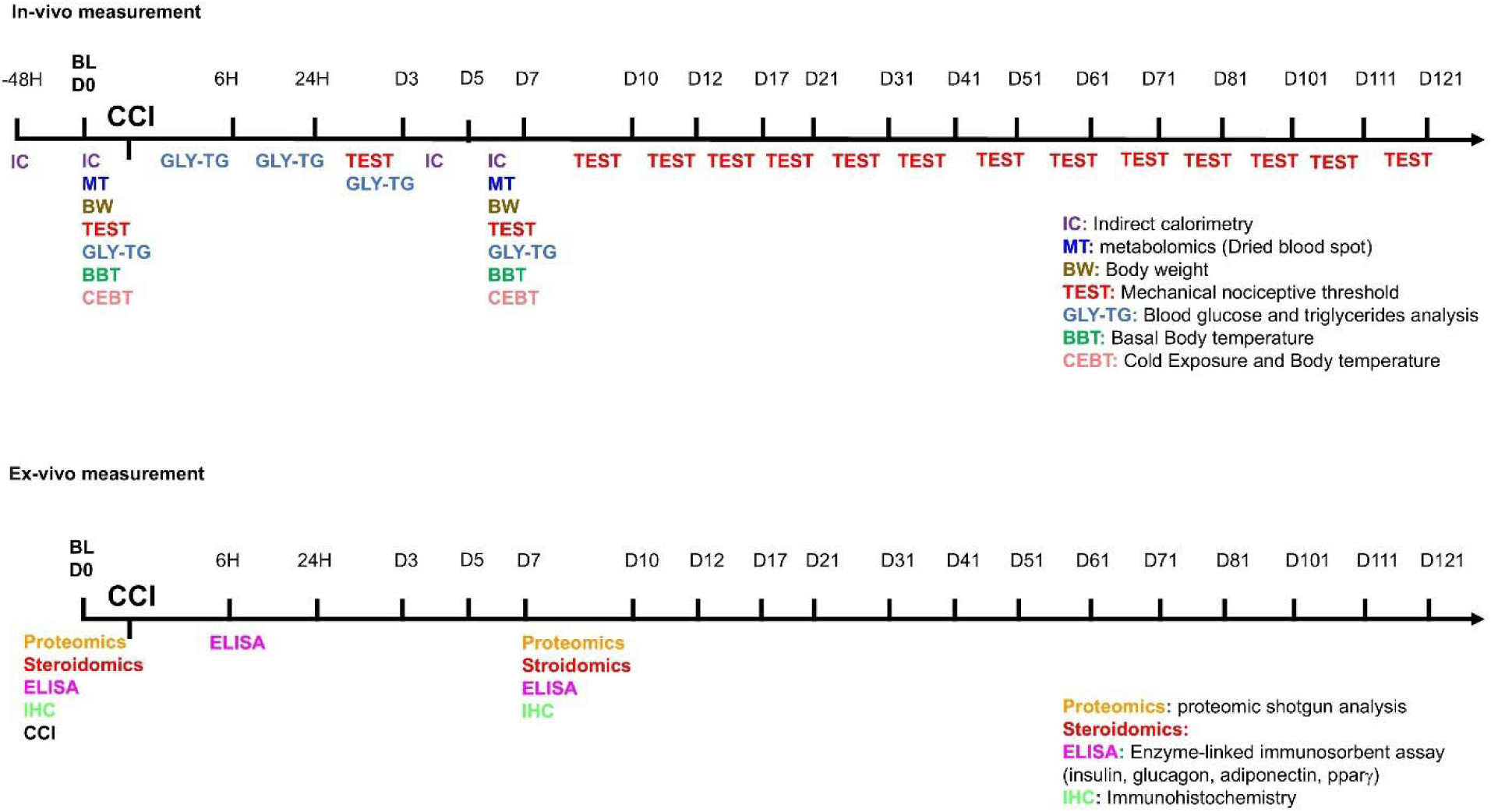
Timeline of experimental procedures. On the line, the hour (6h-24h-48h) or the day of test (D0 to D121) before (baseline - BL) or after CCI are reported. In multicolor acronyms are indicated in-vivo (top panel) or ex-vivo (bottom panel) experiments.

### Assessment of Mechanical Allodynia (Dynamic Plantar Aesthesiometer Test)

Neuropathy onset was evaluated by assessing the sensitivity of both ipsilateral and contralateral hindpaws to normally non-painful mechanical stimuli at various time points, ranging from postoperative day 3 (D3) up to day 121 (D121). To measure nerve injury-induced mechanical allodynia, we employed the Dynamic Plantar Aesthesiometer (Ugo Basile, Model 37400), a device that progressively applies mechanical force over time. The force was directed to the plantar surface of the mouse hindpaw, and the nociceptive threshold was determined as the amount of force (in grams) at which the animal withdrew its paw. On each testing day, the withdrawal response of both ipsilateral and contralateral hindpaws was recorded across three consecutive trials, ensuring at least 10 seconds between each measurement. The final withdrawal threshold was calculated as the average of these three trials. Behavioral testing was conducted in both male and female mice.

### Measurement of Glycemia and Triglycerides

Blood glucose and triglyceride levels were assessed using the Multicare Test Strips system (Biochemical Systems International). Measurements were obtained through tail vein sampling in naïve animals (baseline, BL) and at multiple time points following chronic constriction injury (CCI), specifically at 6 hours, 24 hours, day 3 (D3), and day 7 (D7) [3]. These assessments were conducted in both male and female mice [18].

### Enzyme-Linked Immunosorbent Assay (ELISA) for Insulin, Glucagon, PPAR**γ**, Adiponectin, and Leptin

Serum concentrations of insulin and glucagon were determined using ELISA kits (RayBio® Mouse Insulin ELISA Kit; RayBiotech Inc., Norcross, GA, USA, and Quantikine® ELISA Glucagon Immunoassay; R&D Systems), following previously established protocols^10^. Leptin levels in serum were quantified using the Mouse Leptin (LEP) ELISA Kit (BT LAB) according to the manufacturer’s instructions, and absorbance values were obtained using the Varioskan Lux Instrument (ThermoFisher). Blood samples were collected at 6 hours and 7 days post-CCI (n=3 per time point) from both male and female mice. White adipose tissue (WAT) was dissected from the subcutaneous abdominal (inguinal) adipose panniculus. PPARγ and adiponectin levels in WAT lysates were measured using ELISA kits from MyBiosource Inc. (San Diego, USA) and Quantikine® ELISA Immunoassay (R&D Systems), respectively. Tissue samples were collected from both male and female mice (n=3 per time point) at baseline and seven days post-CCI. All incubation and washing steps were performed according to the manufacturers’ instructions, followed by the addition of a substrate solution. The intensity of the colorimetric reaction was directly proportional to the concentration of insulin and glucagon in serum, as well as PPARγ and adiponectin in adipose tissue. Absorbance was recorded at 450 nm, with wavelength correction at 570 nm for insulin, glucagon, and adiponectin detection.

### Immunohistochemical Analysis

The sciatic nerve from mice in each experimental group was collected for immunofluorescence (IF) analysis. Animals were euthanized using a sub-lethal dose of a Rompun (Bayer) and Zoletil (Virbac) mixture, followed by perfusion with saline and 4% paraformaldehyde in phosphate-buffered saline (PBS, pH 7.4). The sciatic nerve was then extracted and immersed in 4% paraformaldehyde in PBS for 48 hours. Subsequently, the tissue was cryoprotected in a 30% (w/v) sucrose solution in PBS before being stored at -80°C. Cryostat-generated sections (20 µm) were mounted directly onto glass slides for further processing. IF analysis was conducted at baseline (BL) and seven days post-CCI (D7) in both male and female mice (12 months - 12M). For double IF staining, tissue sections were incubated overnight with primary antibodies targeting S100β (Schwann cell marker; mouse monoclonal, 1:100, Sigma-Aldrich: S2532) and IRS1 (Insulin receptor substrate 1; rabbit polyclonal, 1:100, Abcam). Both antibodies were diluted in 0.3% Triton-X100 (Sigma-Aldrich) for optimal penetration.

Following three washes in PBS, sciatic nerve sections were incubated at room temperature for 2 hours with fluorescently labeled secondary antibodies: Alexa Fluor 488-conjugated donkey anti-mouse (1:100, Jackson ImmunoResearch) and Rhodamine-conjugated anti-rabbit (1:100, Jackson ImmunoResearch 1110251). After two additional PBS washes, sections were counterstained with the DNA-binding fluorochrome bisBenzimide (Hoechst, 1:1000, Sigma-Aldrich) for 10 minutes. To attain specificity and eliminate nonspecific secondary antibody signals, negative controls were prepared by staining both control and treated sections with secondary antibodies alone.

### Confocal Imaging and Analysis

Immunostained sections were visualized using laser scanning confocal microscopy (TCS SP5, Leica Microsystems). To prevent spectral overlap and signal bleed-through, images were acquired in sequential scanning mode. High-magnification images (40X) of sciatic nerve sections were processed using I.A.S. software (Delta Systems, Italy). Fluorescence intensity of IRS1 protein was quantified using ImageJ software (version 1.41, National Institutes of Health, USA). At least two sections per nerve were analyzed, with fluorescence quantification performed by converting pixel intensity into brightness values using the RGB (red, green, and blue) method. This approach allows for the digital detection and analysis of biological samples in confocal microscopy [21].

### Body Temperature Measurement

Core body temperature (BT) was recorded rectally using a digital thermometer with a precision of 0.1°C. Measurements were taken under baseline (BL) conditions and after 5 hours of exposure to a cold environment (4°C) across all experimental groups. The same procedure was repeated in the same animals seven days post-injury (D7) for both male and female mice.

### Energy Metabolism Assessment

Energy expenditure (EE), metabolic rate (MR), oxygen consumption (VO₂), and respiratory quotient (RQ) were assessed using an indirect calorimetry (IC) system (TSE PhenoMaster/LabMaster System®) with a constant airflow rate of 0.35 L/min. Mice (N=9–11 per group) were acclimated to the metabolic chamber for 6 hours before data collection. VO₂ was recorded every 20 minutes for individual mice, starting at 7:00 PM and continuing automatically for 48 hours to allow comparison between the dark and light phases of the cycle. Room temperature was maintained at a constant 22°C (±1°C). The respiratory exchange ratio (RER), calculated as the ratio of CO₂ produced to O₂ consumed (RER = VCO₂/VO₂), served as an indicator of substrate utilization. EE was determined using the formula: EE=(3.815+1.232×VCO2/VO2)×VO2EE = (3.815 + 1.232 \times VCO₂/VO₂) \times VO₂EE=(3.815+1.232×VCO2/VO2)×VO2 as provided by the TSE System. EE and RER were analyzed across the entire 48-hour recording period, with specific attention to animals’ resting conditions (i.e., values recorded during activity counts between 0 and 3). Locomotor activity was evaluated during the IC session by tracking the number of infrared beam interruptions. Each cage in the IC system was equipped with the InfraMot® device, which utilizes passive infrared sensors to detect and record mouse movement by capturing body-heat images and monitoring spatial displacement over time. All forms of body movement were detected and recorded as activity data.

### Targeted Metabolomics by Flow Injection Analysis–Tandem Mass Spectrometry (FIA-MS/MS)

Dried blood spot (DBS) samples were collected by spotting whole blood from each mouse onto a filter paper card. The metabolic profile, including 18 amino acids (AAs), free carnitine (C0), and 30 acylcarnitines (ACCs), was analyzed using Flow Injection Analysis–Tandem Mass Spectrometry (FIA-MS/MS) with the NeoBase Non-Derivatized MSMS Kit (PerkinElmer Life and Analytical Sciences, Turku, Finland). ^6, 9,10,13, 43,44^ For metabolite quantification, isotopically labeled internal standards (ISs) were added to each analyte before extraction. Disks of 3.2 mm (corresponding to 3.0–3.2 μL of whole blood) were punched out from DBS samples and quality controls (QCs) and placed into a polypropylene microtiter plate. An extraction solution containing ISs (100 μL) was then added to each well. The ISs, along with the extraction solution and QCs, were provided in the NeoBase Kit (PerkinElmer Life and Analytical Sciences, Turku, Finland). The microplate was sealed and shaken at 700 rpm for 50 minutes at 45°C in a thermo mixer. Subsequently, 75 μL of the supernatant from each well was transferred to a new microplate and loaded into the autosampler for analysis. Both low and high QCs were processed in duplicate under identical conditions as the experimental samples. Metabolic profiling of DBS samples was conducted using a Liquid Chromatography Tandem Quadrupole Mass Spectrometry (LC-MS/MS) system (Alliance HT 2795 HPLC Separation Module coupled to a Quattro Ultima Pt ESI, Waters Corp., Manchester, UK). The mass spectrometer operated in positive electrospray ionization (ESI) mode, employing multiple reaction monitoring (MRM) for data acquisition. MassLynx V4.0 software (Waters Corp.) was used for instrument control, while data processing was automated via NeoLynx (Waters Corp.). Sample injections (30 μL) were introduced directly into the ion source through a narrow PEEK tube, with an injection-to-injection cycle of 1.8 minutes. The ionization source parameters were optimized to maximize ion yield for each metabolite, with the following settings: capillary voltage at 3.25 kV, source temperature at 120°C, desolvation temperature at 350°C, and collision cell gas pressure at 3–3.5 × 10⁻³ mbar (Argon) [18,22–26]

The list of analyzed metabolites and a description of abbreviations as used in the text are available in Table 1

**Table 1.** FIA-MS/MS acquisition parameters used for the determination of whole blood amino acids (AAs) and acylcarnitines (ACCs). MS/MS transitions for each analysed AA and ACC and the corresponding internal standard (IS, shown in bold), the optimal cone potential (V), and collision energy (eV) are shown for each analyte. The capillary potential was 3.5 kV.

| Abbreviation<br>IS | Full name | Transition | Cone<br>potential | Collision<br>energy |
| --- | --- | --- | --- | --- |
| Ala<br>D4Ala | Alanine | 90.2>44.3<br>94.2>48.3 | 40 | 6 |
| Arg<br>His<br>D5Arg | Arginine<br>Histidine | 175.3>70.3<br>156.2>110.2<br>180.3>75.3 | 45 | 17 |
| Cit<br>D2Cit | Citrulline | 176.2>113.2<br>178.2>115.2 | 45 | 13 |
| Gly<br>D2Gly | Glycine | 76.2>30.3<br>78.2>32.3 | 40 | 5 |
| Leu/Ile/Pro-OH<br>Asp<br>Glu<br>Asn<br>D3Leu | Leucine/Isoleucine/Hydroxyproline<br>Aspartic Acid<br>Glutamic Acid<br>Asparagine | 132.2>86.3<br>134.2>88.2<br>148.2>84.2<br>133.2>74.2<br>135.2>89.3 | 40 | 8 |
| Met<br>D3Met | Methionine | 150.2>104.2<br>153.2>107.2 | 45 | 9 |
| Orn<br>Lys/Gln<br>D6Orn | Ornithine<br>Lysine/Glutamine | 133.3>70.3<br>147.2>130.2<br>139.3>76.3 | 40 | 12 |
| Phe<br>D6Phe | Phenylalanine | 166.2>120.2<br>172.2>126.2 | 45 | 11 |
| Tyr<br>D6Tyr | Tyrosine | 182.2>136.2<br>188.2>142.2 | 45 | 12 |
| Val<br>Ser<br>Thr<br>D8Val | Valine<br>Serine<br>Threonine | 118.2>72.3<br>106.1>60.3<br>120.2>74.3<br>126.2>80.3 | 40 | 8 |
| Co<br>D9Co | Free Carnitine | 162.2>103.2<br>171.2>103.2 | 60 | 14 |
| C2<br>D3C2 | Acetylcarnitine | 204.2>85.2<br>207.2>85.2 | 60 | 14 |
| C3 | Propionylcarnitine | 218.2>85.2 | 60 | 15 |
| D<sub>3</sub>C<sub>3</sub> |  | 221.2>85.2 |  |  |
| C<sub>4</sub><br>C<sub>3</sub>DC/C<sub>4</sub>OH<br>D<sub>3</sub>C<sub>4</sub> | Butyrylcarnitine<br>Malonylcarnitine/3-Hydroxy-<br>butyrylcarnitine | 232.3>85.2<br>248.3>85.2<br>235.3>85.2 | 60 | 15 |
| C<sub>5</sub><br>C<sub>5:1</sub><br>C<sub>4</sub>DC/C<sub>5</sub>OH<br>D<sub>9</sub>C<sub>5</sub> | Valerylcarnitine<br>Tiglylcarnitine<br>Methylmalonylcarnitine/3-Hydroxy-valerylcarnitine | 246.2>85.2<br>244.2>85.2<br>262.2>85.2<br>255.2>85.2 | 70 | 16 |
| C<sub>6</sub><br>D<sub>3</sub>C<sub>6</sub> | Hexanoylcarnitine | 260.3>85.2<br>263.3>85.2 | 65 | 16 |

**Table 2.** Concentration levels (ng/mL) for calibrators and QC materials of each steroid monitored in the LC-MS/MS method of analysis are summarized.

| Analytes | Calibration Levels<br>(ng/mL) |  |  |  |  |  |  | QC Levels<br>(ng/mL) |  |  |
| --- | --- | --- | --- | --- | --- | --- | --- | --- | --- | --- |
|  | *L1* | *L2* | *L3* | *L4* | *L5* | *L6* | *L7* | *QC1* | *QC2* | *QC3* |
| CCONE | 0.29 | 0.70 | 1.68 | 4.03 | 10.0 | 24.9 | 62.0 | 0.84 | 4.18 | 31.0 |
| 11-DECOL | 0.08 | 0.20 | 0.49 | 1.17 | 2.91 | 7.23 | 18.0 | 0.24 | 1.21 | 9.0 |
| DHEA | 0.31 | 0.73 | 1.76 | 4.22 | 10.5 | 26.1 | 65.0 | 0.88 | 4.38 | 32.5 |
| DHEAS | 12.9 | 31.0 | 74.4 | 179 | 444 | 1110 | 2750 | 37.2 | 185 | 1380 |
| ADIONE | 0.08 | 0.20 | 0.49 | 1.17 | 2.91 | 7.23 | 18.0 | 0.24 | 1.21 | 9.0 |
| TESTO | 0.03 | 0.08 | 0.20 | 0.47 | 1.16 | 2.89 | 7.20 | 0.10 | 0.49 | 3.60 |
| 17-OHP | 0.12 | 0.29 | 0.70 | 1.69 | 4.20 | 10.5 | 26.0 | 0.35 | 1.75 | 13.0 |
| PROG | 0.12 | 0.29 | 0.69 | 1.72 | 4.27 | 10.6 | 26.5 | 0.36 | 1.78 | 13.2 |

### Label-Free Differential Proteomic Shotgun Analysis

Proteomic analysis was conducted on total proteins extracted from adipose tissue collected from four experimental groups (n=3 mice per pool, with three technical replicates per group). The groups consisted of male and female mice at baseline (BL) and seven days post-injury (D7). White adipose tissue (WAT) was dissected from the subcutaneous abdominal (inguinal) adipose panniculus. Protein concentration was determined, and 50 μg of total protein from each sample was processed using the filter-aided sample preparation (FASP) protocol, which integrates protein purification and enzymatic digestion, as previously described^55^. Samples were transferred onto a Microcon-10 Centrifugal Filter with a 10 kDa molecular weight cut-off (Millipore). After buffer exchange with Urea Buffer (UB: 8M urea, 100mM Tris-HCl, pH 8.5), proteins were denatured, reduced in 8mM dithiothreitol (DTT) for 15 minutes at 56°C, and alkylated with 0.05M iodoacetamide (IAA) for 20 minutes at room temperature. Prior to digestion, the buffer was exchanged with 0.05M ammonium bicarbonate (AmBic). Trypsin digestion was performed at an enzyme-to-protein ratio of 1:50 (w/w) for 16–18 hours at 37°C. The reaction was halted by adding formic acid (FA) to a final concentration of 0.2% (v/v), and peptides were recovered in 0.05M AmBic, concentrated using a speedvac, and stored at -80°C until further analysis. After digestion, peptides were resuspended in 0.1% FA, and 0.300 µg of each sample was spiked with 300 fmol of MassPREP Enolase Digestion Standard (Waters Corp.) as an internal reference. Each sample was analyzed in four technical replicates. Tryptic peptides were separated using an ACQUITY M-Class System (Waters Corp.). A 3 µL aliquot of the digested sample was injected onto a Symmetry C18 5 μm, 180 μm × 20 mm precolumn (Waters Corp.), followed by separation using a 90-minute reversed-phase gradient (2–40% acetonitrile over 75 minutes) at a flow rate of 300 nL/min on an HSS T3 C18 1.8 μm, 75 μm × 150 mm nanoscale LC column (Waters Corp.) maintained at 40°C. Mobile phase A consisted of 0.1% FA in water, while mobile phase B contained 0.1% FA in acetonitrile (ACN). Peptide analysis was performed using a High-Definition Synapt G2-Si Mass Spectrometer (Waters Corp.), coupled directly to the chromatographic system.

Data acquisition followed a data-independent acquisition (DIA) workflow (MSE mode). The instrument was operated with the following settings: electrospray ionization in positive mode (ES+), mass range acquisition of 50–2000 m/z, capillary voltage of 3.2 kV, source temperature of 80°C, cone voltage of 40V, time-of-flight (TOF) resolution of 20,000, precursor ion charge state range of 0.2–4, trap collision energy of 4 eV, transfer collision energy of 2 eV, precursor MS scan time of 0.5 sec, and fragment MS/MS scan time of 1.0 sec. Lock mass correction was performed post-acquisition using the doubly charged monoisotopic ion of [Glu1]-Fibrinopeptide B (Waters Corp.), sampled every 30 seconds. Continuum LC-MS data from three replicate runs per sample were analyzed for qualitative and quantitative assessment using Progenesis QI for Proteomics v4.1 (Nonlinear Dynamics, Waters Corp.). Protein identification was performed by searching against the Mus musculus database (UniProt.Swiss-Prot release 2021_04), with the sequence of Enolase 1 from *Saccharomyces cerevisiae* (UniProtKB/Swiss-Prot AC: P00924) appended for absolute quantification. Search parameters included:

- Trypsin as the specified digestion enzyme
- Automatic tolerance for precursor and product ions
- Minimum of 3 fragment ions matched per peptide
- Minimum of 7 fragment ions matched per protein
- At least 1 peptide matched per protein
- Allowance of 1 missed cleavage
- Carbamidomethylation of cysteine as a fixed modification
- Oxidation of methionine as a variable modification
- False discovery rate (FDR) set at ≤1% at the protein level

For quantitative analysis, the “Absolute Quantitation Using HiN” option in Progenesis QI software was used, with MassPREP Enolase Digestion Standard as an absolute calibrant. The three most abundant peptides per protein (N=3) were measured for quantification [27]. Mass spectrometry proteomics data were deposited in the **ProteomeXchange Consortium** via the **PRIDE** repository with dataset identifier **PXD061152** (DOI: 10.6019/PXD061152).

### Serum Steroid Profiling by UPLC-MS/MS

Corticosterone (CCONE), 11-deoxycortisol (11-DECOL), dehydroepiandrosterone (DHEA), dehydroepiandrosterone sulfate (DHEAS), 4-androstene-3,17-dione (ADIONE), testosterone (TESTO), 17α-hydroxyprogesterone (17-OHP), and progesterone (PROG), along with their corresponding isotopically labeled internal standards (2H8-CCONE, 2H5-11-DECOL, 2H6-DHEAS, 2H5-ADIONE, 2H5-TESTO, 2H8-17-OHP, 2H9-PROG), were obtained from the CHSTM MSMS Steroids Kit (PerkinElmer®, Turku, Finland). Blood samples (approximately 200 μL per mouse) were collected at BL and seven days post-nerve lesion via decapitation. Samples were kept at room temperature (23 ± 1°C) to allow coagulation, then centrifuged at 4°C for 15 minutes at 1400 g. Serum was carefully collected, avoiding the layer immediately above the buffy coat, and transferred into polypropylene tubes. Each endogenous steroid (1 mg) was dissolved in ethanol and stored at -20°C. Stock solutions were diluted in a methanol/water (50:50) solution to obtain a final concentration of 0.03 μM (tuning solution). The internal standard (IS) mix provided in the PerkinElmer® kit was reconstituted in 1.25 mL of acetonitrile (ACN). A daily Precipitation Solution (DPS) containing ISs was prepared by diluting the IS mix 1:100 in ACN with 0.1% FA. Calibrators and quality control (QC) samples, derived from human serum, were obtained from the CHSTM MSMS Steroids Kit (PerkinElmer®, Turku, Finland). The concentration levels (ng/mL) of each steroid analyzed in the LC-MS/MS method are summarized in Table 3.

**Table 3:** MS/MS operating conditions. Multiple reaction monitoring (MRM) functions and settings for detection of steroids are shown. Italics denotes qualifier ion.

| MRM Function | Time Window (min) | Analyte | Transitions (m/z) | Cone Volts | Coll Energy (eV) |
| --- | --- | --- | --- | --- | --- |
| 1 | 5.0–8.0 | CCONE | 346.98 > 121.08 | 38 | 24 |
|  |  | *CCONE* | 346.98 > 97.09 | 38 | 24 |
|  |  | <sup>2</sup>H<sub>8</sub>-CCONE | 354.98 > 125.08 | 38 | 24 |
| 2 | 5.0–8.0 | 11-DECOL | 346.98 > 109.06 | 40 | 28 |
|  |  | *11-DECOL* | 346.98 > 97.09 | 40 | 28 |
|  |  | <sup>2</sup>H<sub>5</sub>-11-DECOL | 351.98 > 100.09 | 40 | 28 |
|  | 4.25–7.0 | DHEAS | 271.2 > 197.1 | 32 | 18 |
|  |  | *DHEAS* | 271.2 > 213.2 | 32 | 18 |
|  |  | <sup>2</sup>H<sub>6</sub>-DHEAS | 277.1 > 219.2 | 32 | 18 |
|  | 6.0–9.5 | DHEA | 271.2 > 197.1 | 32 | 17 |
|  |  | *DHEA* | 271.2 > 213.2 | 32 | 17 |
|  |  | <sup>2</sup>H<sub>8</sub>-OHP |  |  |  |
| 3 | 5.5–8.5 | ADIONE | 287.04 > 97.03 | 36 | 24 |
|  |  | *ADIONE* | 287.04 > 109.06 | 36 | 24 |
|  |  | <sup>2</sup>H<sub>5</sub>-ADIONE | 292.04 > 100.03 | 36 | 24 |
| 4 | 6.0–9.0 | TESTO | 289.04 > 97.09 | 38 | 26 |
|  |  | *TESTO* | 289.04 > 109.05 | 38 | 26 |
|  |  | <sup>2</sup>H<sub>5</sub>-TESTO | 294.04 > 100.09 | 38 | 26 |
| 5 | 7.5–9.5 | 17-OHP | 331.04 > 97.09 | 40 | 32 |
|  |  | *17-OHP* | 331.04 > 109.06 | 40 | 32 |
|  |  | <sup>2</sup>H<sub>8</sub>-OHP | 339.04 > 100.09 | 40 | 32 |
| 7 | 7.5–10.5 | PROG | 315.204 > 97.09 | 40 | 24 |
|  |  | *PROG* | 315.04 > 109.05 | 40 | 24 |
|  |  | <sup>2</sup>H<sub>9</sub>-PROG | 324.204 > 100.09 | 40 | 24 |

### Data Analysis

All results are presented as mean ± SEM. The sample size for in vivo experiments was determined in advance using Power analysis (G*Power 3.1). Statistical analyses were selected based on the nature of the data. Depending on the dataset, comparisons were conducted using either an unpaired t-test, one-way analysis of variance (ANOVA), or two-way repeated-measures ANOVA. For experiments with small sample sizes (N<5 animals) and groups larger than three, non-parametric analysis was performed using the Kruskal-Wallis test. In cases of multiple comparisons, post-hoc analysis was conducted using the Tukey-Kramer test, while t-tests were applied for single comparisons. Statistical significance was set at P < 0.05. Statistical analyses were carried out using Statview 5.0, RStudio, and Python. The effects of sex and CCI on whole blood amino acid (AA) and acylcarnitine (ACC) profiles were assessed using two-way repeated-measures ANOVA, followed by Fisher’s post-hoc test. Sex and CCI were considered independent factors. When data did not meet the assumption of homoscedasticity, ANOVA was performed after Aligned Rank Transformation (ART) for nonparametric factorial analysis. ART allows for nonparametric factorial analyses using standard ANOVA procedures. Statistical significance was considered at P < 0.05. For statistical analysis, the following software was utilized: GraphPad Prism 6.0, Statview 5.0, RStudio, Statistica 6.0 (StatSoft, Tulsa, OK, USA), and MetaboAnalyst.

## RESULTS

### Onset and evolution of neuropathy and pain in aging mice

To evaluate the progression of neuropathic pain (NeP) and allodynic response, we used the model of chronic constriction injury (CCI) of the sciatic nerve[28] and assessed the allodynic response via measurement of pain threshold to mechanical stimulation. The investigation of differences in body weight (BW), estrous cycle status, and baseline mechanical thresholds were performed in male and female mice at three distinct ages - 6 months (6M, adult and fertile period), 12 months (12M, onset of aging), and 18 months (18M, aging phase). BW increased significantly with age, particularly in males (Figure 1A). Vaginal smear analyses of 6M females confirmed the presence of regular estrous cycles. Cytological analyses in 12M females revealed menopause-like changes in 40% of the animals (Supplemental Table 1). Mechanical thresholds at baseline showed no significant differences between males and females, neither at 6M nor at 12M (Figure 1B). Further analysis, including only the 12M females and the estrous cycle status (cycling vs. non-cycling), indicated no significant effect of the hormonal status on mechanical thresholds (Supplemental Table 1). By contrast, at 18M, a reduction in mechanical threshold was observed in females, and thus the presence of significant allodynia (Figure 1B). The study of NeP progression was performed on 12-month-old male and female mice at the onset of senescence and infertility (peri-menopause/menopause in females). Sensitivity and response to painful stimulation were monitored by the Dynamic Plantar

Aesthesiometer test continuously for 120 days, including the aging phase (up to 16 months of age) (Figure 1C). Previous studies[10],[11] have highlighted differences in NeP and allodynic response of adult mice subjected to CCI. In our previous study, adult females (4 months old) showed an initial decrease of NeP compared to males but exhibited a chronicization of pain up to 120 days of duration. In contrast, males showed full recovery within 60 days following peripheral nerve injury. Here, in aged subjects, both the onset and progression of allodynia showed a different pattern of expression. The difference between males and females previously observed in adulthood was not evident in aging mice. Both cohorts exhibited heightened sensitivity to mechanical stimulation beginning on day 3 post-injury, followed by a gradual decline in sensitivity. However, especially around day 70 from the CCI, 12M males showed a higher withdrawal response indicative of persistence of NeP, with a trend towards recovery only after day 100. Conversely, aged females showed a complete recovery from NeP from day 70, showing a subsequent complete resolution of chronic pain. The dynamic of functional recovery in older animals suggests sex-specific differences in nocifensive response. Hence, the time-dependent evolution of recovery is different between aged males and females, and in particular, aged male mice exhibited a marked deterioration in recovery time compared to younger males. This prolonged period of recovery in aged males highlights age-related vulnerabilities in the mechanisms of pain resolution. These data reveal that aging significantly affects the trajectory of recovery from neuropathic pain, with a protracted persistence of NeP in aged males but with sex-specific differences becoming less marked in older cohorts.

### Sex-dependent metabolic changes following neuropathy

The study explored sex-dependent metabolic and thermoregulatory changes in 12-month-old mice subjected to CCI. The findings report significant differences in metabolic responses to NeP between males and females, underscoring age- and sex-specific adaptations. In vivo glycemic levels (Figure 1, panel D) revealed significant sex-specific differences. Aged male mice showed elevated blood glucose levels at 24 hours and day 7 (D7) post-CCI compared to baseline (BL), indicating delayed but prolonged glucose dysregulation. In contrast, females maintained stable glycemic levels across all time points, reflecting resilience to alteration of glucose homeostasis following nerve injury. These findings suggest that male mice are more prone to disruption of glucose homeostasis after CCI. Triglyceride levels (Figure 1, panel E) showed different time points across sexes. Males showed significant reductions in triglyceride levels at 24 hours and day 3 (D3) post-CCI, with levels returning to baseline by D7. Female mice showed consistent reductions at 6 hours, 24 hours, and D3 post-CCI. This pattern suggests enhanced lipid clearance in females, in contrast with the delayed and transitory dysregulation observed in males. Insulin levels revealed sex-specific differences (Figure 1, panel F). In female mice, insulin levels were below the manufacturer’s minimum detectable threshold, suggesting significant age-related decline in insulin levels [3]. Conversely, male mice displayed transient post-CCI hyperinsulinemia at 6 hours. Glucagon analysis further revealed sex differences, with significant differences observed at baseline and 6 hours post-CCI, reflecting distinct regulatory mechanisms of glucose and energy homeostasis between sexes. Thermoregulatory capacity differed between males and females (Figure 1, panel G). Male mice showed a significant pre-CCI decrease in body temperature during cold exposure, with only partial post-CCI recovery at D7. In contrast, female mice maintained stable body temperature, demonstrating adaptive thermoregulatory mechanisms despite aging and NeP. These findings disclosed a decline of thermoregulatory function in aging males, while females appear to maintain a correct thermoregulation.

Insulin receptor substrate 1 (IRS1) expression in the sciatic nerve was analyzed under baseline (naïve) and post-CCI conditions (D7) (Figure 2). Confocal imaging revealed distinct patterns of IRS1 localization co-stained with S100 Calcium Binding Protein B (S100β), a Schwann cell (SC) marker. Fluorescence analysis indicated a significant overexpression of IRS1 following CCI in both sexes, suggesting its upregulation as a possible compensatory response to increased energy demand during nerve repair. This response is consistent with whole-body metabolic changes observed in aged mice, emphasizing the role of insulin signaling in peripheral nerve injury and SC function. The sex-dependent differences in metabolism, including glucose dysregulation, lipid changes, and insulin signaling, demonstrate the complex interplay between aging, sex, and neuropathy. While male mice showed prolonged metabolic and thermoregulatory deficits, female mice showed resilient mechanisms to maintain homeostasis. Upregulation of IRS1 in SCs further supports the role of insulin signaling in nerve regeneration, suggesting potential therapeutic targets to address age- and sex-specific metabolic vulnerabilities in the expression of NeP.

**Figure 2.**
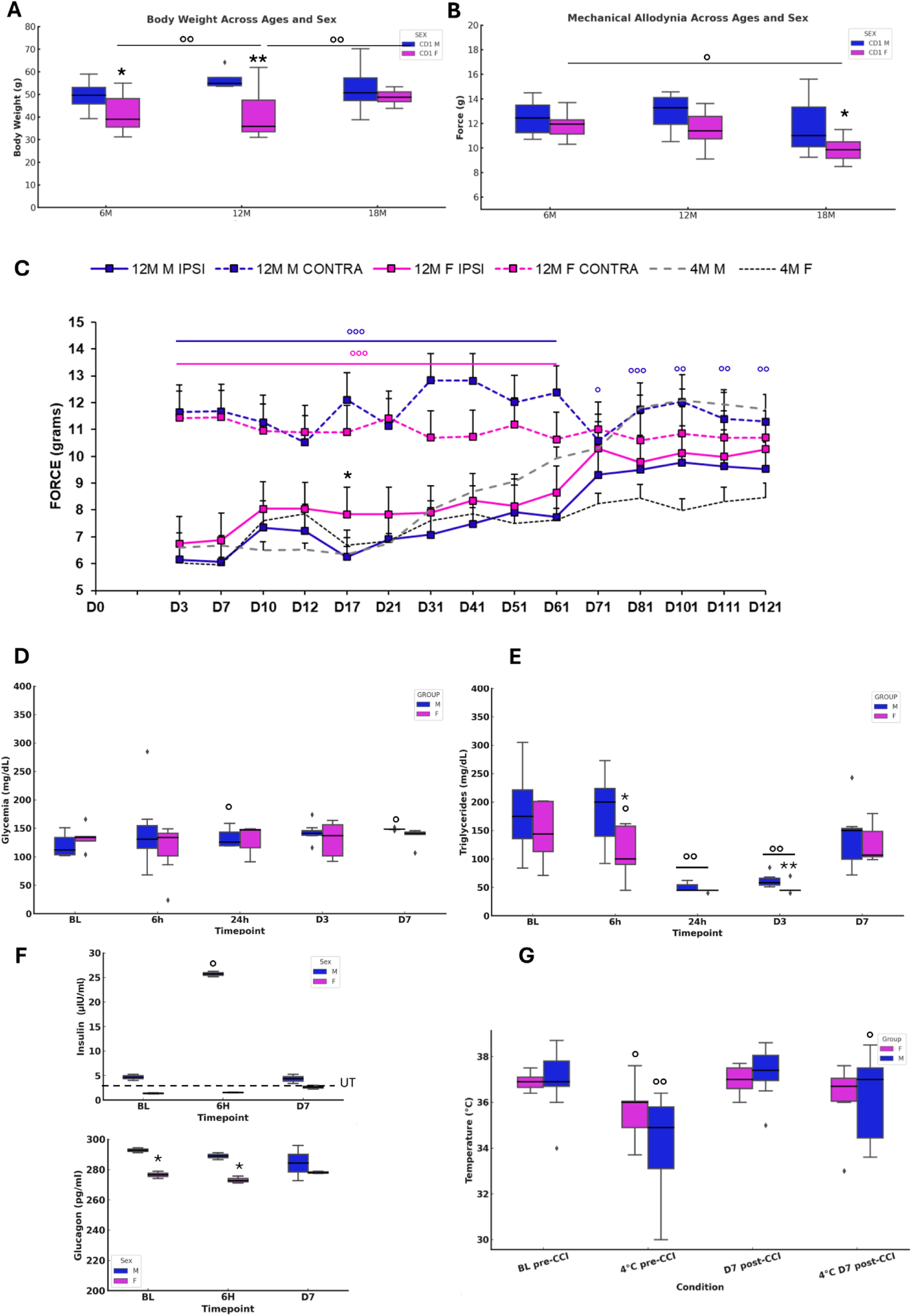
Onset and evolution of Neuropathy and metabolic changes in aging mice. A) Body weight in naïve animals aged 6 months (6M), 12 months (12M), and 18 months (18M). Sample size: N=8–10 per group. ANOVA revealed significant effects for SEX (F₁,₂ = 19.647, p < 0.0001), while AGE (p = 0.0823) and the SEX-AGE interaction (p = 0.0543) were not significant. Tukey-Kramer post-hoc test identified significant differences between: 6M M vs. 6M F (p = 0.0272); 12M M vs. 12M F (p = 0.0008); 6M F vs. 12M F (p = 0.0015); and 12M F vs. 18M F (p = 0.0394). B) Mechanical threshold in naïve animals at 6M, 12M, and 18M. Sample size: N=8–10 per group. ANOVA showed significant effects for SEX (F₁,₂ = 6.569, p = 0.0133) and AGE (F₂,₂ = 4.335, p = 0.0181), while the SEX-AGE interaction was not significant (p = 0.6980). Tukey-Kramer post-hoc test indicated significant differences between: 6M F vs. 18M F (p = 0.0371) and 18M M vs. 18M F (p = 0.0499). * = p < 0.05; ** = p < 0.005 vs. MALE; ° = p < 0.05; °° = p < 0.005 vs. AGE. C) Mechanical allodynia in male and female animals aged 12M at baseline (BL) and following chronic constriction injury (CCI) from 3 days (D3) post-CCI up to 121 days (16M). Sample size: N=11 per group. The dotted lines represent adult mice (4M^55^) data. Repeated measures ANOVA revealed significant effects for SEX (F₃,₄₀ = 371.708, p < 0.0001), TIME (F₄₀,₁₄ = 14.227, p < 0.0001), and SEX-TIME interaction (F₄₂,₅₆₀ = 9.127, p < 0.0001). Unpaired t-test at D17 revealed significant differences in IPSI F vs. IPSI M (t₂₀ = 5.859, p < 0.0001). ° = p < 0.05; °° = p < 0.005; °°° = p < 0.0001 IPSI vs. CONTRA; * = p < 0.05 vs. MALE. D & E) Glucose and triglyceride levels in the blood at baseline (BL) and at 6h, 24h, day 3 (D3), and day 7 (D7) post-CCI in 12M mice (in vivo test). Sample size: N=7 per group. D) Glucose: ANOVA for repeated measures showed no significant differences for SEX, TIME, or SEX-TIME interaction. Paired t-test revealed significant increases compared to BL at 24h (t₆ = -3.34, p = 0.016) and D7 (t₆ = -3.73, p = 0.0097) in males. E) Triglycerides: ANOVA for repeated measures showed a significant effect for TIME (F₁₂,₄ = 29.138, p < 0.0001) but not for SEX or SEX-TIME interaction. Unpaired t-tests showed significant differences between males and females at 6h (t₁₂ = -2.273, P = 0.0422) and D3 (t₁₂ = -2.425, p = 0.0320). Paired t-tests indicated significant decreases in males at 24h (t₆ = 4.54, P = 0.0039) and D3 (t₆ = 4.59, p = 0.0037) and in females at 6h (t₆ = 4.56, p = 0.0039), 24h (t₆ = 5.06, p = 0.0023), and D3 (t₆ = 4.43, p = 0.0044). ° = p < 0.05; °° = p < 0.005 vs. BL; * = p < 0.05, ** = p < 0.005 M vs. F. F) Insulin and glucagon levels in 12M naïve animals (BL) and at 6h and D7 post-CCI. For insulin, female levels were below the manufacturer’s minimum detectable dose and are presented as under threshold (UT) values. This data suggests a dramatic decrease in female insulin levels during aging compared to younger mice [3]. Mann-Whitney U test showed a significant increase in male insulin at 6h post-CCI (p = 0.0495; ° = p < 0.05). For glucagon, the Kruskal-Wallis test (N=3 per group/time-point) revealed significant differences among groups (H₅ = 11.556, p = 0.0408). Mann-Whitney U test showed significant differences between males and females at BL and 6h post-CCI (p<0.05). G) Basal (BL) body temperature and thermoregulation after 4°C exposure pre- and post-CCI in 12M animals. Repeated measures ANOVA showed no SEX differences but a significant effect for cold exposure (pre- and post-CCI; F₂₀,₃ = 16.875, p < 0.0001). Paired t-tests showed a significant decrease in body temperature after 4°C exposure pre-CCI in males (t₆ = 5.331, p = 0.0003) and females (t₆ = 4.019, p = 0.0024). Comparisons of pre-CCI vs. 4°C D7 post-CCI revealed significant increases only in males (t₆ = -2.75, p = 0.020).

### Sex-dependent energy metabolism following neuropathy

By the indirect calorimetry (IC) analysis, we assessed *in vivo* energy expenditure (EE), resting EE (i.e., in lack of motor activity) (REE), and energy substrate oxidation (respiratory exchange ratio, RER) (Figure 3). Continuous calorimetry recording for 48-h in BL condition disclosed several sex-dependent and CCI-dependent differences in energy metabolism. As compared to male mice, female mice showed lower levels of both EE (panel a) and REE (panel b) at BL, while no differences in RER levels were detected. Peripheral nerve injury at D7 was revealed to induce a decrease of EE in the male mice group, as compared to the pre-CCI (i.e., BL) condition (panel a).

**Figure 3.**
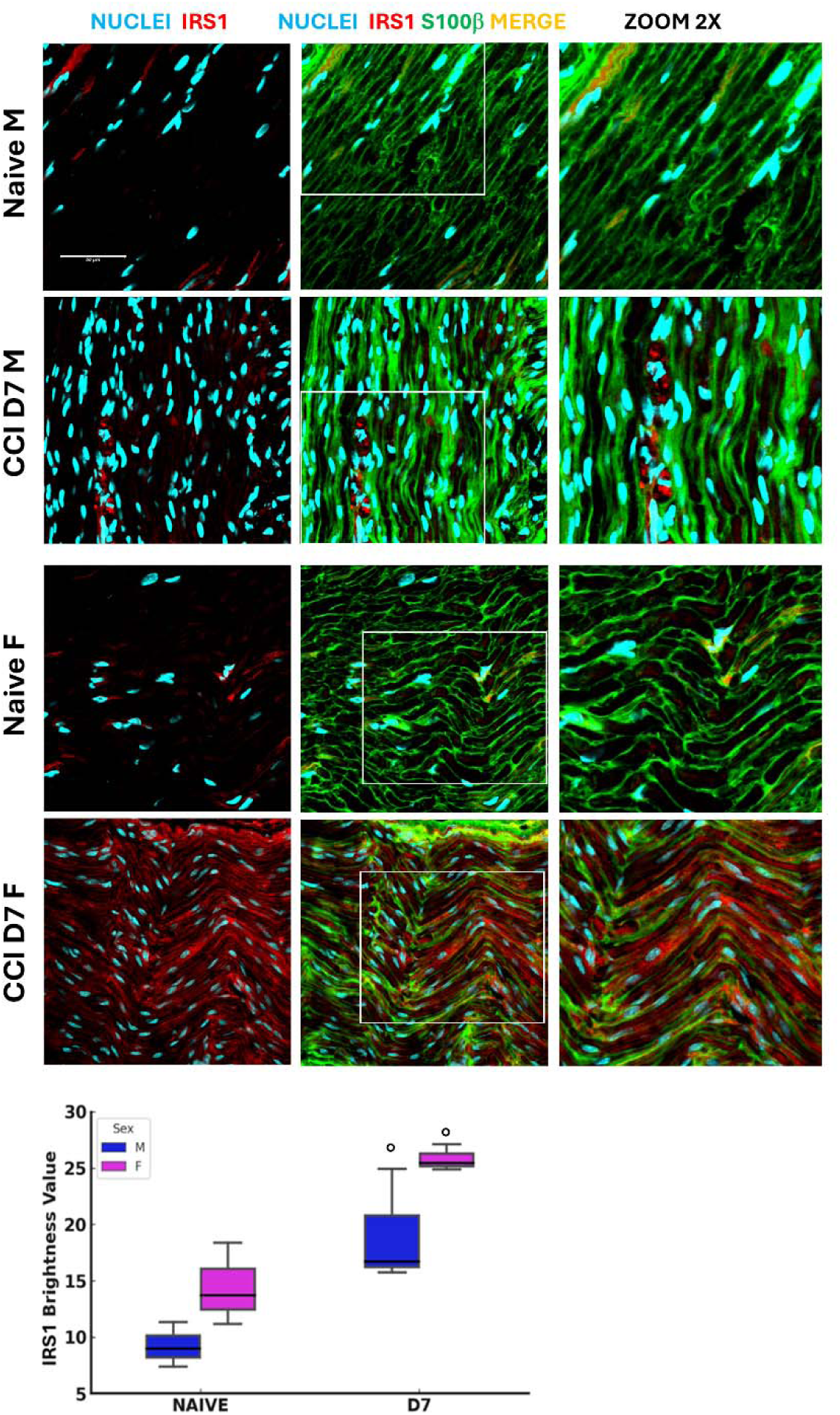
Insulin Receptor Substrate 1 Expression Following Neuropathy. Confocal images (40×1.25x) of sciatic nerve sections co-stained for S100β (green, Schwann cell marker) and IRS1 (red, Insulin receptor substrate 1) illustrate the distribution of these markers in naïve and CCI (day 7 post-injury) male and female mice (detail in zoom 2x). Scale bar: 50 μm. The graphs show IRS1 expression evaluation in male and female mice at BL and post-CCI (D7) conditions using the RGB analysis, which converts pixels into brightness values. Statistical analysis using the Kruskal-Wallis test (H₃ = 8.641, P = 0.0345) revealed significant differences in IRS1 expression. At BL condition vs CCI (○) p < 0.05 (N = 3 per group/time point).

At difference, peripheral nerve injury (i.e., CCI) did not induce changes in EE in female mice, and levels of EE were not different in female mice between BL and D7 conditions (panel a). On the other hand, female mice showed an increase of REE at D7 as compared to the BL condition (panel b), while such an increase was not detected after CCI in male mice (panel b). Lastly, while no differences were found between male and female mice in BL condition, peripheral nerve damage (i.e., CCI) induced an increase of RER in male mice (BL vs D7) while a robust decrease of RER in female mice (BL vs D7). Consequently, male and female mice displayed different RER levels after CCI (D7 male vs D7 female). These data revealed that peripheral nerve damage decreased whole-body EE only in male mice, while REE was increased after peripheral nerve damage but only in female mice that also displayed lower REE at BL. Such an increase of REE after CCI in female mice appears correlated to the decrease of RER observed in the same mice after peripheral nerve damage. Indeed, peripheral nerve damage appeared to affect the prevalent fuel/energy substrate use, which resulted different between male and female mice. After peripheral nerve injury, the energy substrate oxidation (i.e., RER) switched in female mice from a mixed protein/fatty acids oxidation towards a prevalent fatty acids oxidation, thus revealing a major contribution of fat, rather than carbohydrate, to *in vivo* whole-body energy metabolism. Consequently, nerve injury can alter the glucose-fat metabolism homeostasis and, in females, increase lipid oxidation and EE at rest (REE).

### Metabolomic and steroidomic sex-dependent changes following neuropathy

Targeted metabolomic profiling of 12-month-old mice revealed significant sex- and CCI-dependent changes in amino acid (AA) and acylcarnitine (ACC) levels, both at BL and D7 (Figure 4A, Tables S2, S3). At BL, male mice showed lower short-chain ACCs (e.g., C2) levels than females. Following CCI, male mice showed reduced levels of short-chain ACCs (C2, C4, C5OH/C4DC) and longer-chain ACCs (C14, C16) as compared to females. In contrast, male mice exhibited an increase in odd-chain ACCs (C3, C5) at D7, indicating enhanced catabolism of branched-chain amino acids (BCAAs), which may reveal higher energy demand associated with peripheral nerve regeneration. The profile of AA changes also revealed sex- and nerve injury-specific differences. At D7, male mice showed significantly lower levels of citrulline (Cit), lysine/glutamine (Lys/Gln), and glutamate (Glu) compared to females. Notably, Glu, an important excitatory neurotransmitter, was found elevated in male mice post-CCI, suggesting its potential as a biomarker for peripheral nerve injury-induced NeP.

**Figure 4.**
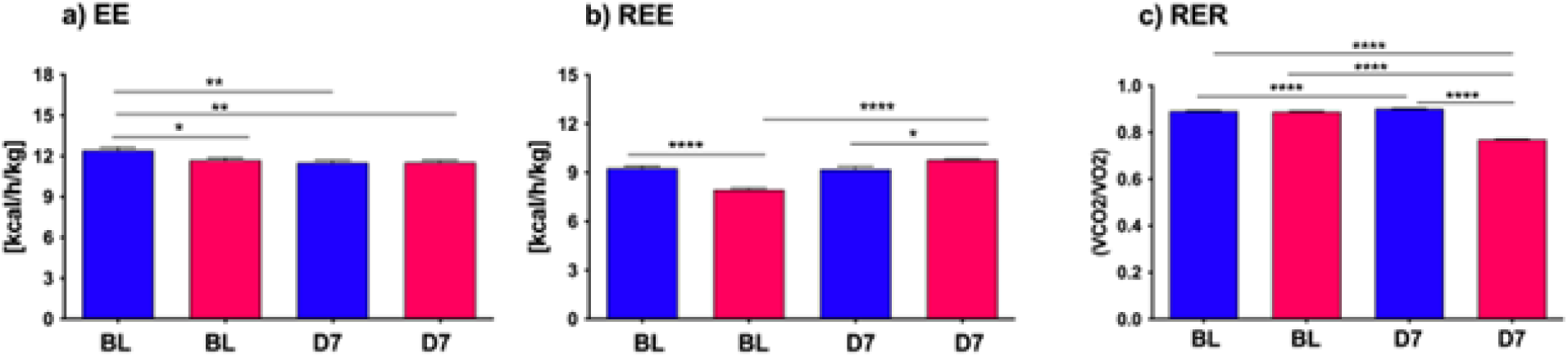
Energy Metabolism in Aging Mice Following Neuropathy. The left panel (a) illustrates the energy expenditure (EE) expressed as kcal/h/kg for both male and female mice at baseline (BL) and day 7 (D7) following chronic constriction injury (CCI). Statistical analysis via two-way ANOVA revealed a significant main effect for CCI/sex factor (F₃,_2921_ = 18.74, p < 0.0001), and a significant main effect for time factor (F_126_,_2921_ = 18.74, p < 0.0001), with further significance between groups observed through Tukey HSD post-hoc testing. The middle panel (b) shows resting energy expenditure (REE), measured as kcal/h/kg, under similar conditions at BL and D7 after CCI. Two-way ANOVA demonstrated a significant main effect for CCI/sex factor (F₃,_276_ = 21.92, p < 0.0001), and no effect of time factor (F_171_,_276_ = 1.23, n.s.); significant differences between groups were further confirmed through Tukey HSD post-hoc analysis. Lastly, the right panel (c) depicts the respiratory exchange ratio (RER), calculated as the VCO₂/VO₂ ratio, for both sexes at BL and D7 post-CCI. Statistical evaluation using two-way ANOVA showed a highly significant main effect for CCI/sex factor (F₃,_2921_ = 18.74, p < 0.0001), a significant main effect for time factor (F_126_,_2921_ = 7.63, p < 0.0001), with differences between groups validated by Tukey HSD post-hoc testing. Statistical significance is indicated as follows: (∗) p < 0.005; (∗∗) p < 0.01; (∗∗∗) p < 0.001; (∗∗∗∗) p < 0.0001.

These findings on aging mice (12 months) expand our earlier work on 4-month-old [3], where we used metabolomics on dried blood spot (DBS) to profile a wide range of metabolites, including AAs and ACCs in male and female mice at baseline and 7 days post-CCI (Tables S2, S3).

Together, these data underscore sex-specific metabolic adaptations after peripheral nerve injury, with male mice showing exclusive patterns of ACC and AA regulation compared to females, particularly in response to the high metabolic and energy demand induced by peripheral nerve injury. Steroid profiling by UPLC-MS/MS further revealed significant sex differences in glucocorticoids and progesterone (PROG) levels in 12-month-old mice at BL and at D7 post-CCI (Figure 4B, Tables 2 and 3). At BL, female mice showed higher levels of corticosterone (CCONE) compared to males, suggesting elevated basal activity of the hypothalamic-pituitary-adrenal (HPA) axis in female mice. After CCI, sex-specific steroid profiles were more evident. Female mice showed a significant increase in CCONE and PROG, indicative of enhanced stress response and potential neuroprotective mechanisms. By contrast, male mice showed significantly lower levels of CCONE and PROG post-injury, suggesting a reduced stress response and impaired metabolic and thermoregulatory adaptation. The increased levels of PROG in females could account for the resilience shown after peripheral nerve injury, as PROG is known for its anti-inflammatory and neuroprotective action. This hormonal effect might contribute to the efficient thermoregulation, as observed in females after CCI. Together, these findings underscore the importance of sex-specific secretion of steroid hormones and the gender differences in the modulation of metabolic responses following damage of peripheral nerve and the development of NeP in aging mice.

### Sex-dependent proteomic changes in adipose tissue following neuropathy

Proteomic profiling of AT revealed distinct sex-specific differences and significant alterations following chronic constriction injury (CCI) in 12-month-old mice. Our experiment was designed to identify the proteins differentially expressed (DEPs) in AT of 12 M-old female and male mice at baseline (BL) or seven days (D7) after sciatic nerve injury [3].

This experiment allowed us the quantification of a set of about 500 protein whose expression is differentially modulated in the four groups of mice (females in baseline condition F_BL; females at day 7 after CCI - F_D7; males in baseline condition - M_BL; males at day 7 after CCI M_D7). From this set we selected 406 DEPs with a maximum fold change (MFC) of the protein expression levels set as MFC ≥1.5 in statistically significant observation (ANOVA, p-value≤0.05). Details on analysis are reported in the supplementary material (Table S6). A heatmap representation (Figure 5) of the 406DEPs dataset, based on protein recurrence, illustrates the different expression regulation between females and males, which results of particular relevance between BL and D7 (post-injury) in both sexes.

**Figure 5.**
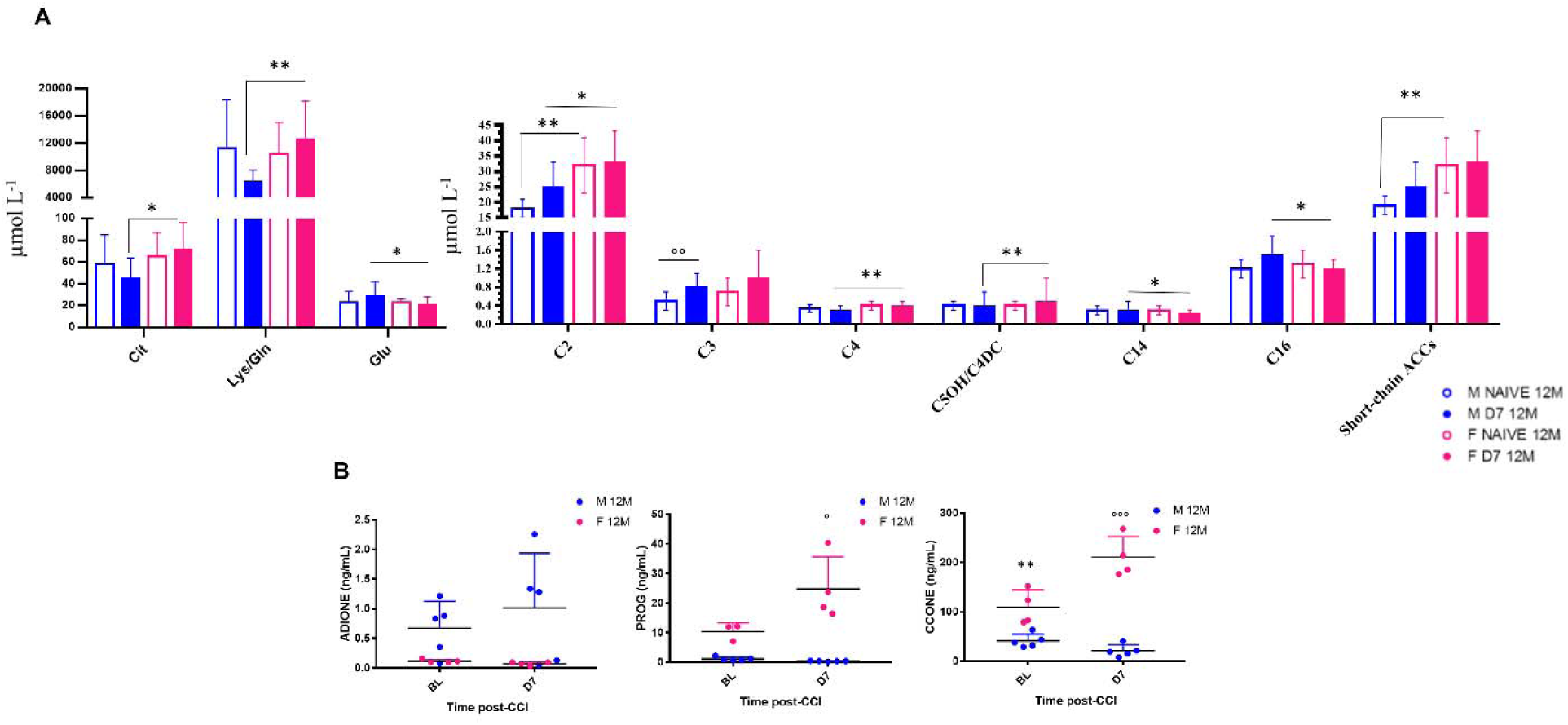
Sex-dependent Metabolomic and Steroidomic changes in aging mice following neuropathy. Bar graphs represent levels of amino acids (AAs) and acylcarnitines (ACCs) expressed as µmol L^−1^ in 12-month-old male and female mice at baseline (Naïve 12M) and 7 days post-CCI (D7 12M). Table of statistics are reported in S2, and S3. Asterisks (*) indicate significant differences between males and females, while circles (°) denote significant differences between baseline and post-CCI conditions. Statistical significance assessed by two-way ANOVA followed by Fischer’s least significance difference (LSD) test is indicated as follows: *p < 0.050, **p < 0.010, ***p < 0.001; °p < 0.050, °°p < 0.010, °°°p < 0.001 (N = 7 per group).B) Scatter plots depict the levels of Androstenedione (4-Androstene-7,13-dione) (ADIONE), progesterone (PROG), and corticosterone (CCONE), expressed in ng/mL, in 12-month-old male (blue dots) and female (pink dots) mice at baseline (BL) and 7 days post-CCI (D7). Asterisks (*) indicate significant differences between males and females at baseline, while circles (°) highlight significant differences between baseline and post-CCI conditions (N = 4–5 per group) (CCONE: p < 0.0001; Tukey’s post-hoc test p = 0.01 for BL males vs. BL females; (Tukey’s post-hoc test p < 0.0001 for BL females vs. D7 females; Tukey’s post-hoc test p < 0.0001 for D7 males vs. D7 females; PROG: Tukey’s post-hoc test p = 0.017 for BL females vs. D7 females; Tukey’s post-hoc test p < 0.0001 for D7 males vs. D7 females

**Figure 6.**
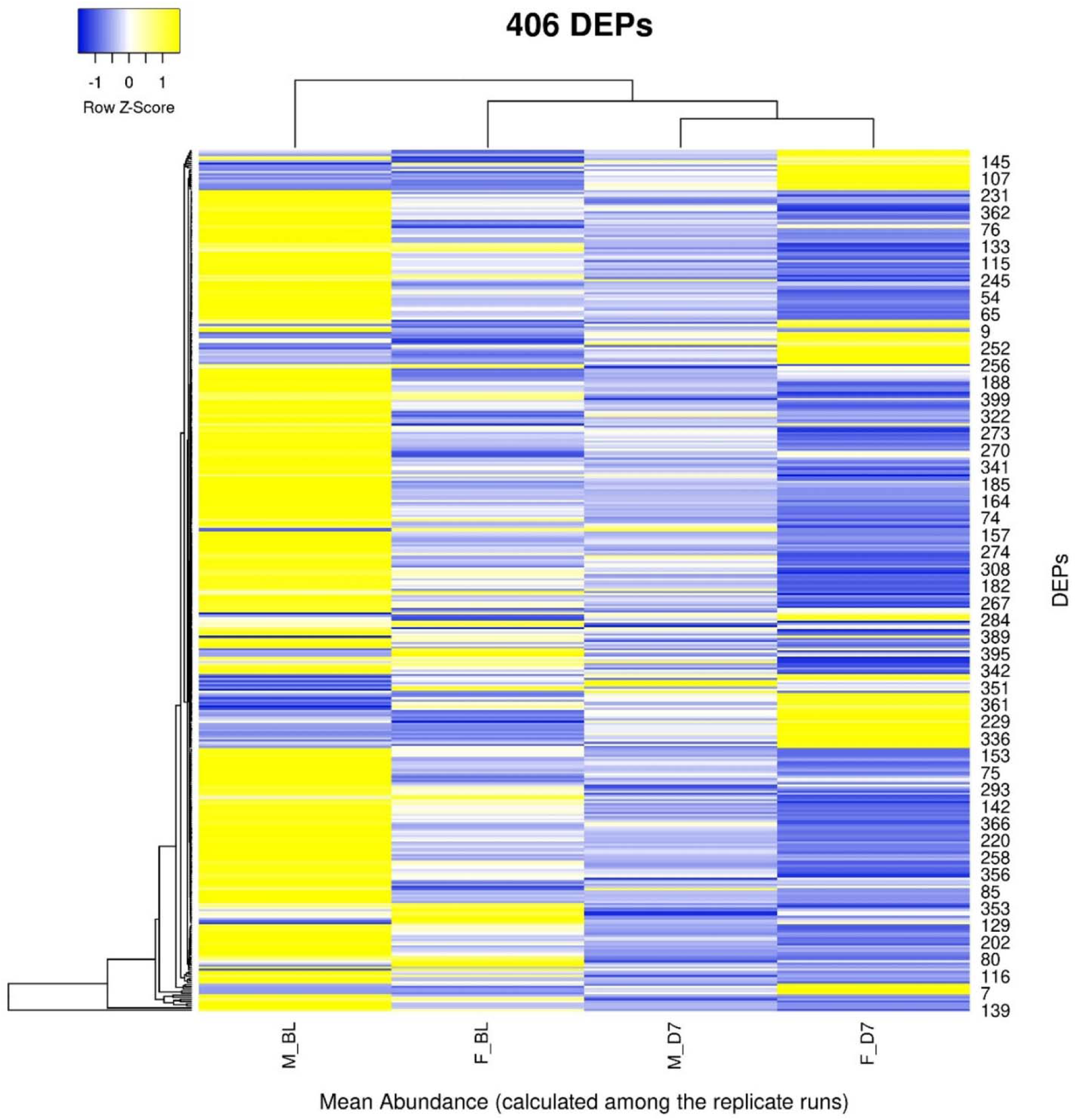
Heatmap representation of the 406 differentially expressed proteins (DEPs) in the four groups. (Complete DEPs list in Table S4) Rows represent the mean abundance of each protein (measured in three replicate runs in LC-MS) in the four groups.

**Figure 7.**
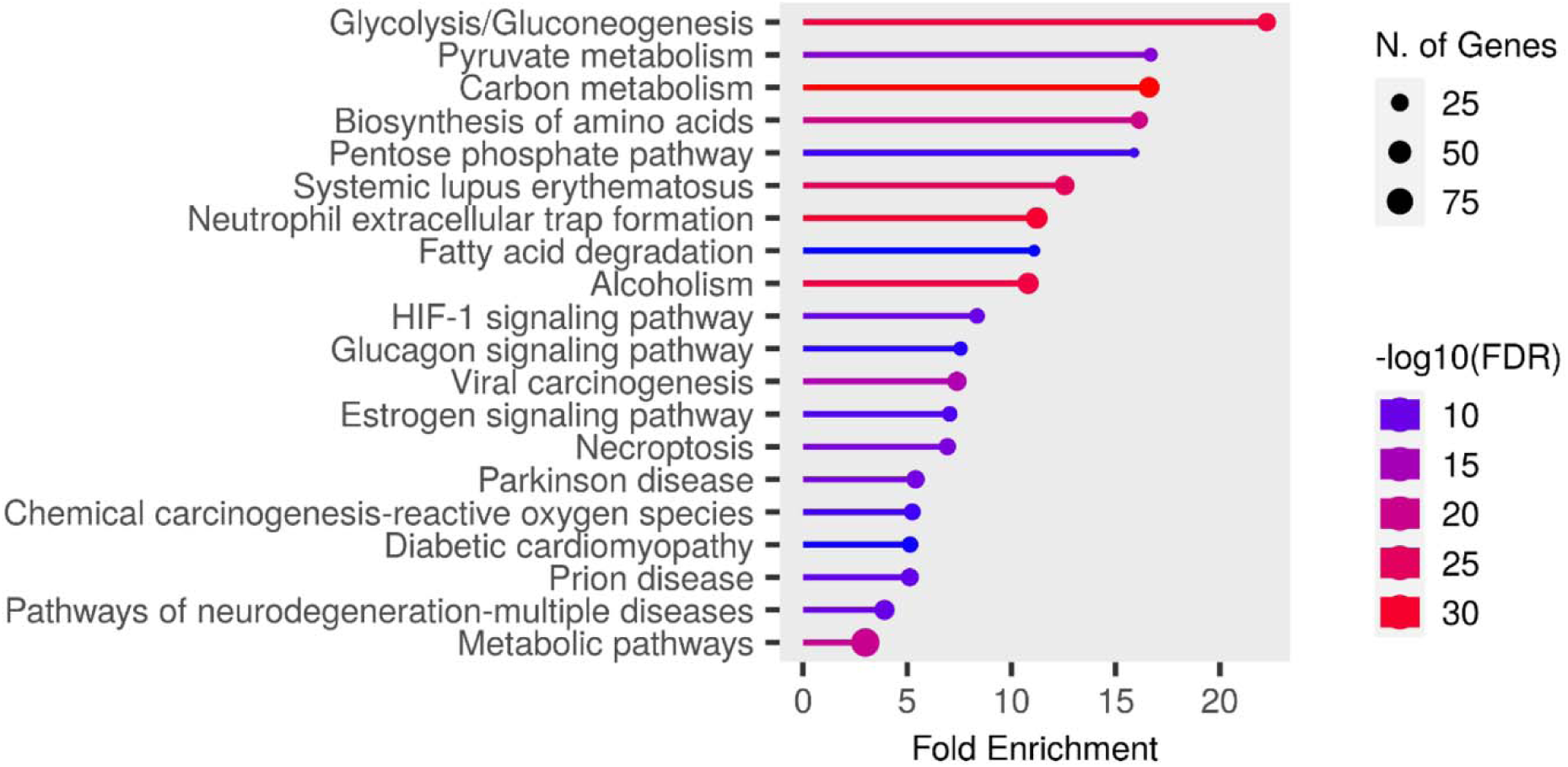
Chart representing enrichment pathway analysis in KEGG pathway database. The chart was obtained using ShinyGO 0.80 (http://bioinformatics.sdstate.edu/go/)

We then focused on sex-related differences in the response to nerve injury by performing an expression analysis using the Reactome database, a platform that allows visualization of expression data mapped onto biological pathway diagrams [23]. To explore these differences, we analyzed the lists of differentially expressed proteins (DEPs) and their fold changes from the comparisons F_BL vs. M_BL and F_D7 vs. M_D7 (Table S6), using males as the baseline reference. The fold change values were used to color-code the elements within the pathway diagrams. As shown in Figure 8, the Voronoi diagram representing all enriched pathways in our datasets highlights those proteins involved in key biological processes, such as Metabolism, Immune System, and Metabolism of Proteins, display opposite regulatory patterns within the same pathways depending on sex. A clear example of this contrasting regulation is observed in the Gluconeogenesis pathway (R-HSA-70263). At baseline, this pathway is primarily enriched with proteins showing higher average expression in males (protein accession numbers: P06745, Q64467, P16858, Q9Z2V4, P21550). However, following nerve injury, the same pathway is instead enriched with proteins exhibiting higher average expression in females (P17751, Q9Z2V4, P09411, P05064, Q91Y97, P05063, P16858, P09041, P21550, O70250). Overall, this analysis indicates that, at baseline, males show greater enrichment of pathways related to lipid metabolism, such as mitochondrial fatty acid oxidation. In contrast, after CCI, females display higher expression of proteins involved in gluconeogenesis (Figures S2 and S3).

**Figure 8.**
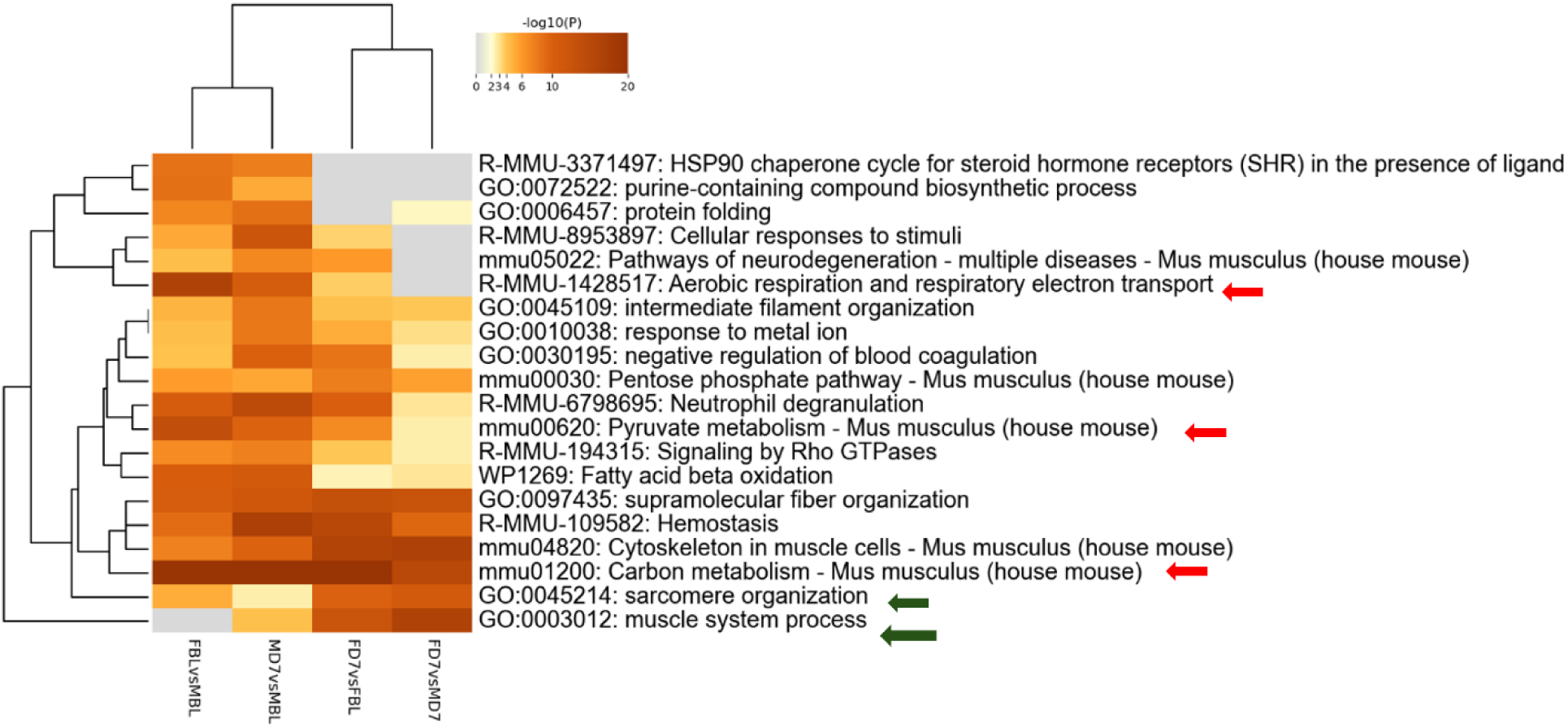
Metascape analysis: enriched terms clusters Statistically. enriched terms (GO/KEGG terms, Reactome canonical pathways, WiKi Pathways) are depicted in the heat map. The term with the best p-value within each cluster as its representative term are displayed in the dendrogram. The heatmap cells are colored by their p-values, white cells indicate the lack of enrichment for that term in the corresponding gene list.

Additionally, in females, we observe increased expression of proteins associated with the Muscle Contraction pathway (R-HSA-397014) following nerve injury—proteins that are not overexpressed in females at baseline (Q5XKE0, P97457, P13412, O09165, P58771, P20801, Q60605, P05977, Q64518, Q8R429, O55143, Q9JI91). This finding may suggest a recovery of muscle strength and activity in the post-injury phase.

### Sex-dependent Adipokines and pPAR***γ*** Changes Following Neuropathy

The analysis of circulating levels of leptin showed no significant differences between sexes or between BL and post-CCI D7, indicating that leptin does not play a major role in the sex-specific metabolic response to neuropathy (Figure S4). In contrast, adiponectin levels showed significant sex- and CCI-dependent changes. In male mice, adiponectin levels decreased significantly after CCI (Figure 9 panel A), suggesting impaired metabolic regulation and a reduced role of adiponectin signaling in anti-inflammatory activity following nerve injury. In contrast, female mice showed a significant post-CCI increase in adiponectin levels, indicating a potential compensatory mechanism to counteract inflammation and metabolic disruption caused by neuropathy. At baseline, male mice had higher adiponectin levels compared to females, supporting the hypothesis that adiponectin plays a more prominent role in male metabolic homeostasis under physiological conditions. However, following CCI, female mice showed significantly higher adiponectin levels than males, revealing a sex-dependent shift in metabolic and hormonal regulation after peripheral nerve injury. This suggests that females may have an enhanced adaptive response to neuropathy, possibly promoting resilience against metabolic stress. Moreover, different sex-dependent patterns were also observed in pPARγ levels (Figure 9 panel B). In both sexes, pPARγ levels in AT were significantly decreased following CCI, indicating potential dysregulation of pPARγ signaling. The reduction was more pronounced in male mice than in females, suggesting a greater male susceptibility to metabolic imbalance after neuropathy, potentially impairing adipose tissue function. Comparisons at BL revealed significantly higher pPARγ levels in male mice than in female mice, reinforcing the idea that pPARγ plays a more central role in male adipose tissue homeostasis. After CCI, this sex difference was even more pronounced, suggesting that nerve injury amplifies pre-existing sex differences and makes males more vulnerable to metabolic dysfunction linked to pPARγ loss.

**Figure 9.**
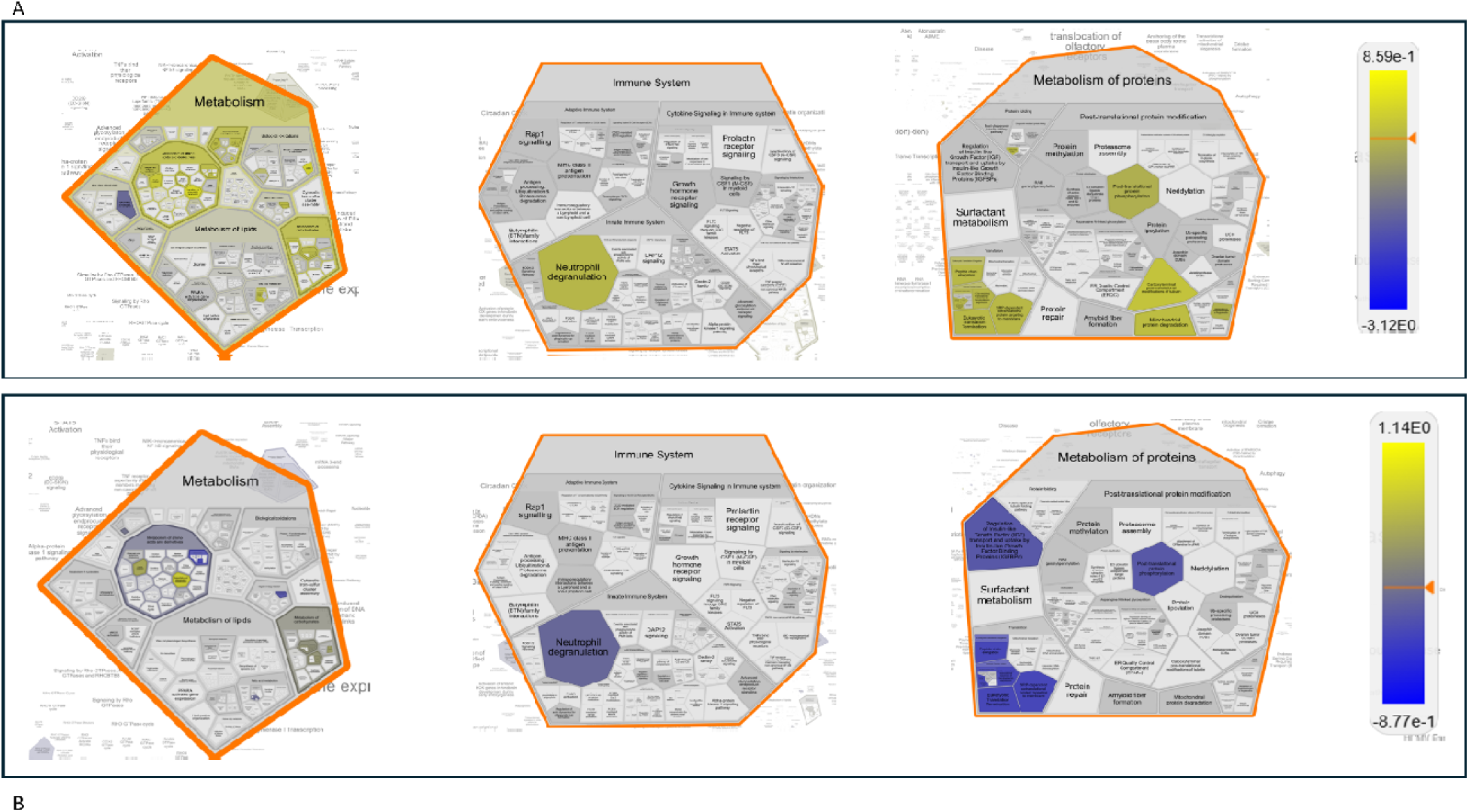
Reactome expression analysis. Voronoi Plot representation of pathways enriched from the differentially expressed proteins (DEPs) with their fold changes in the pairing F_BL vs M_BL (A) and F_D7vs M_D7 (B). Protein expression levels in males where always considered as baseline value of expression. The numeric values are used to color objects in pathway diagrams. Yellow indicates positive value of the Log10_FC, hence protein with a higher expression value in female. Blue indicates negative value of the Log10_FC, hence protein with a higher expression value in male The same pathways are more enriched in female at baseline and in males upon nerve injury (post –CCI).

**Figure 10.**
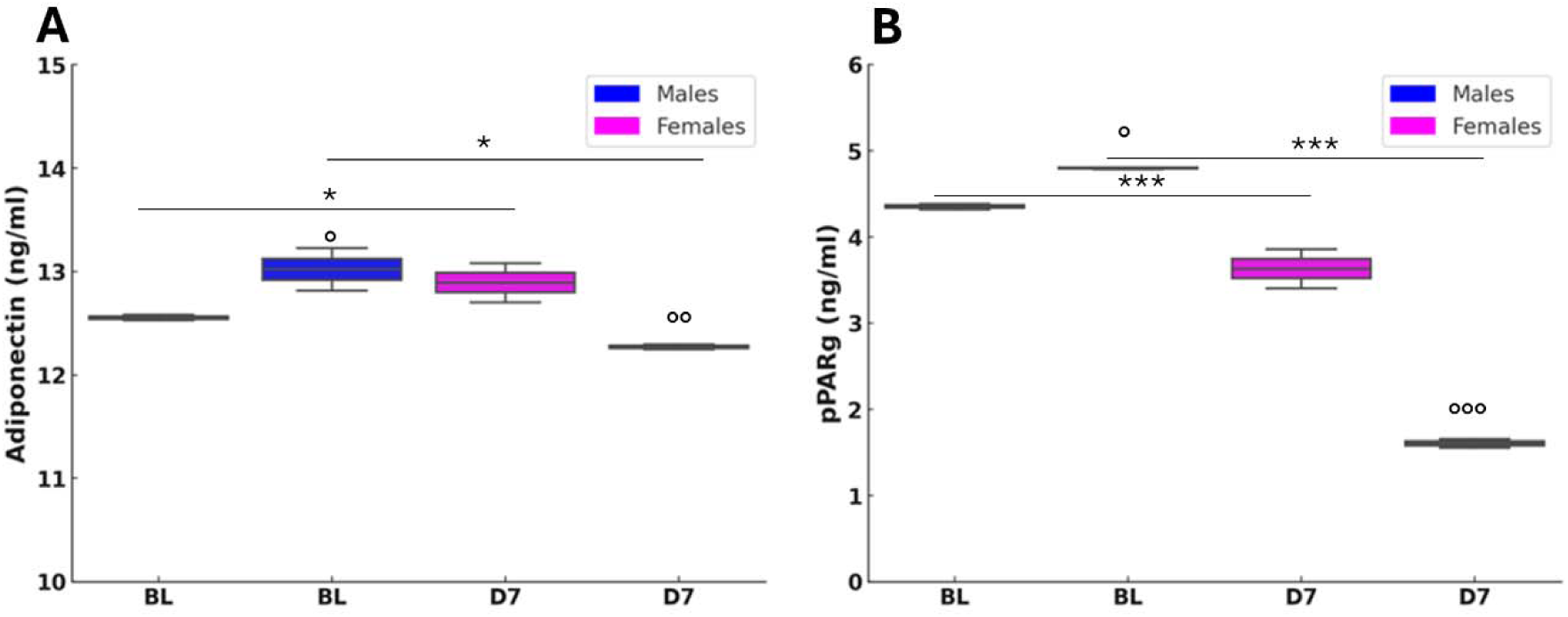
Adiponectin and Peroxisome proliferator-activated receptor gamma (pPARγ) variation after nerve lesion in AT. A) Adiponectin measured by enzyme-linked immunosorbent assay (ELISA) in AT at baseline condition (BL) and 7 days after CCI (D7) in male and female mice. Unpaired t-test: BL M vs D7 M t_4_= 3,879, p = 0.0179 (*); BL F vs D7 F t_4_ = 5,573, p = 0.0051 (*); BL M vs BL F t_4_ = 3,037, p = 0.0385 (°); D7 M vs D7 F t_4_ = -6,219, p = 0.0034 (°°); n=3 group. B) pPARγ measured by ELISA in AT at baseline condition (BL) and 7 days after CCI (D7) in male and female mice. Unpaired t-test: BL M vs D7 M t_4_= 22,629, p <0.0001 (***); BL F vs D7 F t_4_= 15.071, p=0.0001 (**); BL M vs BL F t_4_= -5,452, p= 0.0055 (°); D7 M vs D7 F t4= -96,779, p < 0.0001 (°°°); n=3 group

## DISCUSSION

Our study explored sex-specific differences in neuropathy development and chronic pain persistence during aging, with a focus on whole-body and AT metabolism and their contribution to pain processing and recovery following peripheral nerve injury (CCI). While mechanical thresholds and allodynia sensitivity were comparable between sexes at 6 and 12 months, 18-month-old females showed increased sensitivity, suggesting that aging, rather than hormonal fluctuations (in lack of estrous cycle effects) amplifies pain vulnerability in females. Longitudinal monitoring after CCI revealed that females gradually recovered, achieving full resolution by day 70, whereas males displayed sustained hypersensitivity throughout the 121-day observation period. The delayed recovery in males correlated with pronounced metabolic and thermoregulatory deficits. Hyperglycemia was observed only in males at 24h and 7 days post-CCI, indicating disrupted glucose regulation. Cold exposure before and after CCI revealed that males had lower baseline body temperatures and defective thermoregulatory responses, further deteriorating in post-injury conditions. Conversely, females maintained stable thermoregulation.

The calorimetric analysis provided novel insights into sex-specific energy metabolism following CCI. Females, despite having lower baseline energy expenditure, maintained stable caloric output post-CCI, with a notable increase in resting energy expenditure. This contrasted with the decline observed in males. Additionally, CCI induced a sex-dependent shift in energy substrate use: males slightly increased carbohydrate reliance, while females showed a marked shift toward lipid oxidation, partly explaining their increased energy expenditure. Insulin has a neuroprotective effect on nerve regeneration, and the IR-1-mediated signaling may modulate peripheral nerve injury recovery. The study of diabetic neuropathy in rats has shown that the deficit in IR signaling is associated with exacerbated hyperalgesia [3] or that intrathecal insulin delivery can accelerate axon regeneration and rescue the retrograde loss of collateral axons after peripheral nerve injury [32,33] Compared to pre-CCI levels, insulin receptor-1 (IR-1) expression increased in male and female mice following experimental neuropathy. These findings suggest that the upregulation of IR-1 is a key mechanism supporting nerve regeneration and recovery from peripheral nerve damage in both sexes, not only in diabetic neuropathy models but also in non-diabetic neuropathy induced by traumatic nerve injury.

Targeted metabolomic profiling revealed robust sex- and CCI-dependent differences in amino acid (AA) and acylcarnitine (ACC) metabolism, highlighting distinct mitochondrial responses to nerve injury. At baseline, female mice showed higher levels of short-chain ACCs, particularly acetylcarnitine (C2), compared to males, suggesting more active fatty acid turnover under physiological conditions. Following CCI, both sexes displayed an overall increase in circulating ACCs, but females maintained significantly elevated C2 levels, along with a marked increase in butyrylcarnitine (C4). In contrast, only males showed an elevation in propionylcarnitine (C3), indicating divergent branched-chain amino acid (BCAA) catabolism between sexes [34,35]. Furthermore, while CCI did not significantly alter long-chain ACCs (e.g., C14, C16) in males, these metabolites were markedly reduced in females. This pattern suggests enhanced mitochondrial fatty acid oxidation in females post-injury, consistent with the observed decrease of respiratory exchange ratio (RER) and shift toward lipid utilization. The reduction in C14 and C16 long-chain ACCs in females may reflect more efficient flux through β-oxidation, potentially supported by increased activity of very-long-chain acyl-CoA dehydrogenase (VLCAD), leading to greater conversion into acetyl-CoA and subsequent entry into the tricarboxylic acid (TCA) cycle. The metabolic shift observed in female mice following CCI, despite stable dietary intake and nutritional status, is reminiscent of a fasting-like state, characterized by increased lipid oxidation, mitochondrial efficiency, and possibly ketogenesis through acetyl-CoA redirection. This adaptation appears protective, as the accumulation of long-chain ACCs like palmitoylcarnitine, is associated with impaired mitochondrial respiration, membrane hyperpolarization, reactive oxygen species (ROS) generation, and mitochondrial damage [36,37]. The reduction of long-chain ACCs in females likely reflects a beneficial mitochondrial response contributing to neuropathy resolution. This metabolic profile may be shaped by sex-specific regulation of *pPAR*γ and adiponectin in adipose tissue (AT). Following CCI, males showed increased expression of both markers, while females exhibited a reduction [38]. *pPAR*γ, modulated by nutritional status [38,39], may be downregulated in response to sustained fatty acid oxidation in females to preserve homeostasis. This mirrors a caloric restriction (CR)-like state, previously associated with improved neuropathy recovery and autophagy activation in wild-type and Ambra1⁺/gt mice [18,40]. Notably, CR-induced reductions in long-chain ACCs in Ambra1 males resembled those observed in aged females after CCI, paralleling their superior functional recovery. The role of adiponectin remains complex: while it exerts anti-inflammatory effects in metabolic disorders [41], it is also associated with increased pain in inflammatory diseases [42–44], Its reduction in females may support higher energy expenditure and thermogenesis [45–48] Additionally, progesterone, elevated in females post-CCI, has neuroprotective and promyelinating effects via metabolites like allopregnanolone [49–52]. These hormonal changes may contribute to female resilience, as male mice showed reduced corticosterone and progesterone levels post-injury.

Targeted proteomics of AT revealed 406 differentially expressed proteins (DEPs), with CCI-modulating pathways linked to energy metabolism and muscle function. While pathways related to carbon metabolism, aerobic respiration, respiratory electron transport, and pyruvate metabolism were less represented in females than in males post-CCI, pathways associated with sarcomere organization, muscle system processes, and muscle regeneration were significantly enriched in female mice. Interrogation of the Reactome database disclosed the existence of the gluconeogenesis pathway as particularly enriched by proteins such as beta-enolase (*Eno3*), glyceraldehyde-3-phosphate dehydrogenase (*Gapdh*), glucose-6-phosphate isomerase (*Gpi*) and phosphoenolpyruvate carboxykinase (*Pck1*) that were upregulated only before the CCI. Thus, while males showed early enrichment in gluconeogenesis-related proteins, females displayed post-CCI upregulation of glycolytic and pentose phosphate pathway (PPP) components (e.g., GAPDH, ALDOA, PGK1/2, ENO3, TPI1), which support NADPH production, redox balance, and muscle metabolism [53,54]. This metabolic shift may contribute to the prevention of pentose phosphate pathway (PPP) dysregulation induced by polyol pathway activation under hyperglycemic conditions [55–57], a transient state observed exclusively in males and known to interfere with SCs function

[55,58]. In contrast, the upregulation of enzymes such as GAPDH, PGK1/PGK2, and ENO3 can improve glycolysis efficiency [56], reducing fructose accumulation and preventing excessive ROS production [57,59]. In turn, Aldoa (fructose-bisphosphate aldolase A) can contribute to fructose clearance through glycolysis [56], and GAPDH and PGK enzymes help to regulate NAD+/NADH ratios, potentially offsetting the redox imbalance caused by excessive fructose metabolism [57,60]. Moreover, if TPI1 (triosephosphate isomerase) and PGAM2 (phosphoglycerate mutase 2) are upregulated, fructose-derived intermediates are more efficiently redirected into ATP production rather than accumulating in alternative pathways like the polyol pathway. Moreover, PGK1, PGK2, and GAPDH can regulate glycolysis and impact PPP flux through metabolic rewiring. Upon exposure of cells to oxidative challenge, the mRNA levels of enzymes such as GAPDH can increase and undergo oxidation, leading to **t**he diversion of glucose metabolism towards the PPP to generate NADPH and counteract oxidative stress [54,59,60].

The ability of female mice to switch between metabolic states likely reflects enhanced metabolic flexibility. Elevated short-chain ACCs, such as C4, combined with reduced RER, indicate preferential lipid utilization and mitochondrial oxidation, simulating exercise or CR-induced adaptations. This profile enables the redirection of glucose intermediates through protective pathways, limiting ROS accumulation and maintaining redox homeostasis. The convergence of molecular, behavioral, and endocrine findings mirrors human epidemiological trends, where midlife males show increased NeP prevalence and metabolic dysfunction [61]. In contrast, aged females in our study exhibited improved recovery, driven by efficient lipid oxidation, lower long-chain ACC accumulation, downregulation of the polyol pathway, and sustained thermogenic capacity [61]. These data highlight the central role of sex-dependent metabolic adaptations in determining susceptibility and recovery from NeP. Key regulators such as *pPAR*γ, acylcarnitines, insulin signaling, and steroid hormones emerge as potential therapeutic targets. Notably, *pPAR*γ agonists [62] have shown promise in modulating inflammation in surgical pain models, underscoring the translational relevance of these pathways. Altogether, our findings support the development of biomarker-guided, sex- and age-tailored strategies to treat neuropathic pain in aging populations.

### Conclusion: a functional divergence portrayed by metabolic flexibility

Aged females exhibit enhanced metabolic flexibility, defined by increased lipid oxidation, polyol pathway suppression, and improved redox homeostasis, enabling complete NeP recovery. These findings reveal key age- and sex-dependent metabolic adaptations in NeP, with metabolic resilience explaining the enhanced recovery observed in aged females compared to aged males. Peripheral nerve injury imposes substantial metabolic demands, requiring energy-intensive reparative processes. In response, aged females initiate an adaptive metabolic program resembling a fasting state, characterized by preferential lipid utilization, antioxidant defense via the pentose phosphate pathway, neuroprotective steroid biosynthesis, and downregulation of inflammatory and neurotoxic pathways (e.g., the hyper pathway). Conversely, aged males display metabolic inflexibility, accumulation of lipid intermediates, and increased vulnerability to oxidative stress and inflammation. This metabolic divergence between resilience and susceptibility underscores the necessity of age- and sex-specific therapeutic strategies targeting metabolic pathways to enhance neuroprotection and recovery.

## ACKNOWLEDGMENTS

The Italian Ministry of Health GR-2011-02346912; NUTRAGE, Consiglio Nazionale delle Ricerche FOE-2021 DBA.AD005.225

## Data Availability Statement

The mass spectrometry proteomics data have been deposited in the ProteomeXchange Consortium via the PRIDE repository under the dataset identifier PXD061152 (DOI: 10.6019/PXD061152). Additional data and materials are available upon request. Requests for datasets, protocols, or reagents should be directed to the corresponding authors.

## AUTHOR CONTRIBUTIONS

Conceptualization, SM; RC; methodology, SM, CR, LP, GG, IC; Investigation, SM, LP, VV, FDA, CP, VM, GG, IC; writing—original draft, SM, RC; writing—review & editing, SM, RC, CR, ZNY, DC, LP; funding acquisition, SM; resources SM; statistical analysis, SM, CR, DC; supervision, SM; RC.

## DECLARATION OF GENERATIVE AI AND AI-ASSISTED TECHNOLOGIES

During the preparation of this work, the author(s) used ChatGPT 4o to improve readability and language of the work and to generate script for phyton to create graphs or cartoons useful in schematic representations. After using this tool or service, the author(s) reviewed and edited the content as needed and take(s) full responsibility for the content of the publication.

## REFERENCES

[1] Giovannini, S., Coraci, D., Brau, F., Galluzzo, V., Loreti, C., Caliandro, P., Padua, L., Maccauro, G., Biscotti, L., and Bernabei, R. (2021). Neuropathic pain in the elderly. Diagnostics (Basel). 2021 Mar 30;11(4):613. doi: 10.3390/diagnostics11040613.

[2] Bronge, W., Lindholm, B., Elmståhl, S., and Siennicki-Lantz, A. (2024). Epidemiology and Functional Impact of Early Peripheral Neuropathy Signs in Older Adults from a General Population. Gerontology 70. 10.1159/000535620.

[3] Vacca, V., Rossi, C., Pieroni, L., De Angelis, F., Giacovazzo, G., Cicalini, I., Ciavardelli, D., Pavone, F., Coccurello, R., and Marinelli, S. (2023). Sex-specific adipose tissue’s dynamic role in metabolic and inflammatory response following peripheral nerve injury. iScience 26. 10.1016/j.isci.2023.107914.

[4] Katal, S., Taubman, K., Han, J., and Gholamrezanezhad, A. (2023). Aging Muscles, Myositis, Pain, and Peripheral Neuropathies: PET Manifestations in the Elderly. Preprint, 10.1016/j.cpet.2022.09.009

[5] Palmer AK, Jensen MD. METABOLIC CHANGES IN AGING HUMANS: CURRENT EVIDENCE AND THERAPEUTIC STRATEGIES. J CLIN INVEST. 2022 AUG 15;132(16):E158451. DOI: 10.1172/JCI158451.

[6] Quirós Cognuck, S., Reis, W.L., Silva, M., Debarba, L.K., Mecawi, A.S., de Paula, F.J.A., Rodrigues Franci, C., Elias, L.L.K., and Antunes-Rodrigues, J. (2020). Sex differences in body composition, metabolism-related hormones, and energy homeostasis during aging in Wistar rats. Physiol Rep 8. 10.14814/phy2.14597.

[7] Ana, A.P., Pedro, P.H., Frihling, B.E.F., Souza E Silva, P., De Moraes, L.F.R.N., and Migliolo, L. (2023). Adipose tissue, systematic inflammation, and neurodegenerative diseases. Preprint, 10.4103/1673-5374.343891 10.4103/1673-5374.343891.

[8] Numakawa T, Matsumoto T, Numakawa Y, Richards M, Yamawaki S, Kunugi H. PROTECTIVE ACTION OF NEUROTROPHIC FACTORS AND ESTROGEN AGAINST OXIDATIVE STRESS-MEDIATED NEURODEGENERATION. J TOXICOL. 2011;2011:405194. DOI: 10.1155/2011/405194.

[9] Vacca, V., Marinelli, S., Pieroni, L., Urbani, A., Luvisetto, S., and Pavone, F. (2016). 17Beta-estradiol counteracts neuropathic pain: A behavioural, immunohistochemical, and proteomic investigation on sex-related differences in mice. Sci Rep 6. 10.1038/srep18980.

[10] De Angelis, F., Vacca, V., Tofanicchio, J., Strimpakos, G., Giacovazzo, G., Pavone, F., Coccurello, R., and Marinelli, S. (2022). Sex Differences in Neuropathy: The Paradigmatic Case of MetFormin. Int J Mol Sci 23. 10.3390/ijms232314503.

[11] Vacca, V., Marinelli, S., De Angelis, F., Angelini, D.F., Piras, E., Battistini, L., Pavone, F., and Coccurello, R. (2021). Sexually dimorphic immune and neuroimmune changes following peripheral nerve injury in mice: novel insights for gender medicine. Int J Mol Sci 22. 10.3390/ijms22094397.

[12] Vacca, V., Marinelli, S., Pieroni, L., Urbani, A., Luvisetto, S., and Pavone, F. (2014). Higher pain perception and lack of recovery from neuropathic pain in females: A behavioural, immunohistochemical, and proteomic investigation on sex-related differences in mice. Pain 155. 10.1016/j.pain.2013.10.027.

[13] Blaszkiewicz, M., Willows, J.W., Dubois, A.L., Waible, S., DiBello, K., Lyons, L.L., Johnson, C.P., Paradie, E., Banks, N., Motyl, K., et al. (2019). Neuropathy and neural plasticity in the subcutaneous white adipose depot. PLoS One 14. 10.1371/journal.pone.0221766.

[14] Willows, J.W., Robinson, M., Alshahal, Z., Morrison, S.K., Mishra, G., Cyr, H., Blaszkiewicz, M., Gunsch, G., DiPietro, S., Paradie, E., et al. (2023). Age-related changes to adipose tissue and peripheral neuropathy in genetically diverse HET3 mice differ by sex and are not mitigated by rapamycin longevity treatment. Aging Cell 22. 10.1111/acel.13784.

[15] Willows, J.W., Gunsch, G., Paradie, E., Blaszkiewicz, M., Tonniges, J.R., Pino, M.F., Smith, S.R., Sparks, L.M., and Townsend, K.L. (2023). Schwann cells contribute to demyelinating diabetic neuropathy and nerve terminal structures in white adipose tissue. iScience 26. 10.1016/j.isci.2023.106189.

[16] De Angelis, F., Vacca, V., Pavone, F., and Marinelli, S. (2020). Impact of caloric restriction on peripheral nerve injury-induced neuropathic pain during ageing in mice. European Journal of Pain (United Kingdom) 24. 10.1002/EJP.1493.

[17] Marinelli, S., Eleuteri, C., Vacca, V., Strimpakos, G., Mattei, E., Severini, C., Pavone, F., and Luvisetto, S. (2015). Effects of age-related loss of P/Q-type calcium channels in a mice model of peripheral nerve injury. Neurobiol Aging 36. 10.1016/j.neurobiolaging.2014.07.025.

[18] Coccurello, R., Nazio, F., Rossi, C., De Angelis, F., Vacca, V., Giacovazzo, G., Procacci, P., Magnaghi, V., Ciavardelli, D., and Marinelli, S. (2018). Effects of caloric restriction on neuropathic pain, peripheral nerve degeneration and inflammation in normometabolic and autophagy defective prediabetic Ambra1 mice. PLoS One 13. 10.1371/journal.pone.0208596.

[19] Marinelli, S., Nazio, F., Tinari, A., Ciarlo, L., D’Amelio, M., Pieroni, L., Vacca, V., Urbani, A., Cecconi, F., Malorni, W., et al. (2014). Schwann cell autophagy counteracts the onset and chronification of neuropathic pain. Pain 155. 10.1016/j.pain.2013.09.013.

[20] Marinelli, S., Vacca, V., Angelis, F. De, Pieroni, L., Orsini, T., Parisi, C., Soligo, M., Protto, V., Manni, L., Guerrieri, R., et al. (2019). Innovative mouse model mimicking human-like features of spinal cord injury: efficacy of Docosahexaenoic acid on acute and chronic phases. Sci Rep 9. 10.1038/s41598-019-45037-x.

[21] Inman, C. F. et al. Validation of computer-assisted, pixel-based analysis of multiple-colour immunofluorescence histology. J Immunol Methods 302, 156–67 (2005).

[22] Bonomini M, Di Liberato L, Del Rosso G, Stingone A, Marinangeli G, Consoli A, Bertoli S, De Vecchi A, Bosi E, Russo R, Corciulo R, Gesualdo L, Giorgino F, Cerasoli P, Di Castelnuovo A, Monaco MP, Shockley T, Rossi C, Arduini A. Effect of an L-carnitine-containing peritoneal dialysate on insulin sensitivity in patients treated with CAPD: a 4-month, prospective, multicenter randomized trial. Am J Kidney Dis. 2013;62(5):929–38.

[23] Cicalini I, Rossi C, Pieragostino D, Agnifili L, Mastropasqua L, di Ioia M, De Luca G, Onofrj M, Federici L, Del Boccio P. Integrated Lipidomics and Metabolomics Analysis of Tears in Multiple Sclerosis: An Insight into Diagnostic Potential of Lacrimal Fluid. Int J Mol Sci. 2019; 13;20(6):1265.

[24] Di Liberato L, Arduini A, Rossi C, Di Castelnuovo A, Posari C, Sacchetta P, Urbani A, Bonomini M. L-Carnitine status in end-stage renal disease patients on automated peritoneal dialysis. J Nephrol. 2014;27(6):699–706.

[25] Rossi C, Cicalini I, Zucchelli M, di Ioia M, Onofrj M, Federici L, Del Boccio P, Pieragostino D. Metabolomic Signature in Sera of Multiple Sclerosis Patients during Pregnancy. Int J Mol Sci. 2018 Nov 14;19(11):3589.

[26] Rossi C, Marzano V, Consalvo A, Zucchelli M, Levi Mortera S, Casagrande V, Mavilio M, Sacchetta P, Federici M, Menghini R, Urbani A, Ciavardelli D. Proteomic and metabolomic characterization of streptozotocin-induced diabetic nephropathy in TIMP3-deficient mice. Acta Diabetol. 2018;55(2):121–129.

[27] Marini F, Carregari VC, Greco V, Ronci M, Iavarone F, Persichilli S, Castagnola M, Urbani A, Pieroni L. Exploring the HeLa Dark Mitochondrial Proteome. Front Cell Dev Biol. 2020 Mar 5;8:137.

[28] Bennett, G.J., and Xie, Y.K. (1988). A peripheral mononeuropathy in rat that produces disorders of pain sensation like those seen in man. Pain 33. 10.1016/0304-3959(88)90209-6.

[29] Ge, S.X., Jung, D., Jung, D., and Yao, R. (2020). ShinyGO: A graphical gene-set enrichment tool for animals and plants. Bioinformatics 36. 10.1093/bioinformatics/btz931.

[29] Goedhart, J., and Luijsterburg, M.S. (2020). VolcaNoseR is a web app for creating, exploring, labeling and sharing volcano plots. Sci Rep 10. 10.1038/S41598-020-76603-3.

[30] Zhou, Y., Zhou, B., Pache, L., Chang, M., Khodabakhshi, A.H., Tanaseichuk, O., Benner, C., and Chanda, S.K. (2019). Metascape provides a biologist-oriented resource for the analysis of systems-level datasets. Nat Commun 10. 10.1038/S41467-019-09234-6.

[31] Milacic, M., Beavers, D., Conley, P., Gong, C., Gillespie, M., Griss, J., Haw, R., Jassal, B., Matthews, L., May, B., et al. (2024). The Reactome Pathway Knowledgebase 2024. Nucleic Acids Res 52. 10.1093/nar/gkad1025.

[32] Sugimoto, K., Rashid, I.B., Shoji, M., Suda, T., and Yasujima, M. (2008). Early Changes in Insulin Receptor Signaling and Pain Sensation in Streptozotocin-Induced Diabetic Neuropathy in Rats. Journal of Pain 9. 10.1016/j.jpain.2007.10.016.

[33] Toth, C., Brussee, V., Martinez, J.A., McDonald, D., Cunningham, F.A., and Zochodne, D.W. (2006). Rescue and regeneration of injured peripheral nerve axons by intrathecal insulin. Neuroscience 139. 10.1016/j.neuroscience.2005.11.065.

[34] Koves, T.R., Ussher, J.R., Noland, R.C., Slentz, D., Mosedale, M., Ilkayeva, O., Bain, J., Stevens, R., Dyck, J.R.B., Newgard, C.B., et al. (2008). Mitochondrial Overload and Incomplete Fatty Acid Oxidation Contribute to Skeletal Muscle Insulin Resistance. Cell Metab 7. 10.1016/J.CMET.2007.10.013.

[35] McCoin, C.S., Knotts, T.A., and Adams, S.H. (2015). Acylcarnitines-old actors auditioning for new roles in metabolic physiology. Preprint, 10.1038/nrendo.2015.129 10.1038/nrendo.2015.129.

[36] Liepinsh, E., Makrecka-Kuka, M., Volska, K., Kuka, J., Makarova, E., Antone, U., Sevostjanovs, E., Vilskersts, R., Strods, A., Tars, K., et al. (2016). Long-chain acylcarnitines determine ischaemia/reperfusion-induced damage in heart mitochondria. Biochemical Journal 473. 10.1042/BCJ20160164.

[37] Rutkowsky, J.M., Knotts, T.A., Ono-Moore, K.D., McCoin, C.S., Huang, S., Schneider, D., Singh, S., Adams, S.H., and Hwang, D.H. (2014). Acylcarnitines activate proinflammatory signaling pathways. Am J Physiol Endocrinol Metab 306. 10.1152/ajpendo.00656.2013.

[38] Ahmadian, M., Suh, J.M., Hah, N., Liddle, C., Atkins, A.R., Downes, M., and Evans, R.M. (2013). Pparγ signaling and metabolism: The good, the bad and the future. Nat Med 19. 10.1038/nm.3159.

[39] Vidal-Puig, A., Jimenez-Liñan, M., Lowell, B.B., Hamann, A., Hu, E., Spiegelman, B., Flier, J.S., and Moller, D.E. (1996). Regulation of PPAR γ gene expression by nutrition and obesity in rodents. Journal of Clinical Investigation 97. 10.1172/JCI118703.

[40] Hatori, M., Vollmers, C., Zarrinpar, A., DiTacchio, L., Bushong, E.A., Gill, S., Leblanc, M., Chaix, A., Joens, M., Fitzpatrick, J.A.J., et al. (2012). Time-restricted feeding without reducing caloric intake prevents metabolic diseases in mice fed a high-fat diet. Cell Metab 15. 10.1016/J.CMET.2012.04.019.

[41] Fantuzzi, G. (2008). Adiponectin and inflammation: Consensus and controversy. Journal of Allergy and Clinical Immunology 121. 10.1016/j.jaci.2007.10.018

[42] Choi, H.M., Doss, H.M., and Kim, K.S. (2020). Multifaceted physiological roles of adiponectin in inflammation and diseases. Preprint, 10.3390/ijms21041219 10.3390/ijms21041219

[43] Iannitti, T., Graham, A., and Dolan, S. (2015). Adiponectin-mediated analgesia and anti-inflammatory effects in rat. PLoS One 10. 10.1371/journal.pone.0136819

[44] Sun, L., Li, H., Tai, L.W., Gu, P., and Cheung, C.W. (2018). Adiponectin regulates thermal nociception in a mouse model of neuropathic pain. Br J Anaesth 120. 10.1016/J.BJA.2018.01.016

[45] Askari, A., Arasteh, P., Homayounfar, R., Naghizadeh, M.M., Ehrampoush, E., Mousavi, S.M., and Alipoor, R. (2020). The role of adipose tissue secretion in the creation and pain level in osteoarthritis. Endocr Regul 54. 10.2478/enr-2020-0002

[46] Qiao, L., Yoo, H.S., Bosco, C., Lee, B., Feng, G.S., Schaack, J., Chi, N.W., and Shao, J. (2014). Adiponectin reduces thermogenesis by inhibiting brown adipose tissue activation in mice. Diabetologia 57. 10.1007/s00125-014-3180-5

[47] Wang, L., Luo, Y., Luo, L., Wu, D., Ding, X., Zheng, H., Wu, H., Liu, B., Yang, X., Silva, F., et al. (2021). Adiponectin restrains ILC2 activation by AMPK-mediated feedback inhibition of IL-33 signaling. Journal of Experimental Medicine 218. 10.1084/JEM.20191054

[48] Wei, Q., Lee, J.H., Wang, H., Bongmba, O.Y.N., Wu, C.S., Pradhan, G., Sun, Z., Chew, L., Bajaj, M., Chan, L., et al. (2017). Adiponectin is required for maintaining normal body temperature in a cold environment. BMC Physiol 17. 10.1186/s12899-017-0034-7

[49] Koenig, H.L., Schumacher, M., Ferzaz, B., Do Thi, A.N., Ressouches, A., Guennoun, R., Jung-Testas, I., Robel, P., Akwa, Y., and Baulieu, E.E. (1995). Progesterone synthesis and myelin formation by Schwann cells. Science (1979) 268. 10.1126/science.7770777

[50] Patte-Mensah, C., Meyer, L., Taleb, O., and Mensah-Nyagan, A.G. (2014). Potential role of allopregnanolone for a safe and effective therapy of neuropathic pain. Preprint, 10.1016/j.pneurobio.2013.07.004

[51] Schumacher, M., Guennoun, R., Robert, F., Carelli, C., Gago, N., Ghoumari, A., Gonzalez Deniselle, M.C., Gonzalez, S.L., Ibanez, C., Labombarda, F., et al. (2004). Local synthesis and dual actions of progesterone in the nervous system: Neuroprotection and myelination. 10.1016/j.ghir.2004.03.007.

[52] Schumacher, M., Guennoun, R., Mercier, G., Désarnaud, F., Lacor, P., Bénavides, J., Ferzaz, B., Robert, F., and Baulieu, E.E. (2001). Progesterone synthesis and myelin formation in peripheral nerves. In Brain Research Reviews 10.1016/S0165-0173(01)00139-4

[54] Lunt, S.Y., and Vander Heiden, M.G. (2011). Aerobic glycolysis: Meeting the metabolic requirements of cell proliferation. Annu Rev Cell Dev Biol 27. 10.1146/annurev-cellbio-092910-154237.

[54] Xiao, W., Wang, R.S., Handy, D.E., and Loscalzo, J. (2018). NAD(H) and NADP(H) Redox Couples and Cellular Energy Metabolism. Preprint, 10.1089/ars.2017.7216 10.1089/ars.2017.7216.

[55] Hao, W., Tashiro, S., Hasegawa, T., Sato, Y., Kobayashi, T., Tando, T., Katsuyama, E., Fujie, A., Watanabe, R., Morita, M., et al. (2015). Hyperglycemia promotes Schwann cell de-differentiation and de-myelination via sorbitol accumulation and Igf1 protein down-regulation. Journal of Biological Chemistry 290. 10.1074/jbc.M114.631291.

[56] Hers, H.G. (1956). The mechanism of the transformation of glucose in fructose in the seminal vesicles. Biochim Biophys Acta 22. 10.1016/0006-3002(56)90247-5.

[57] Talwar, D., Miller, C.G., Grossmann, J., Szyrwiel, L., Schwecke, T., Demichev, V., Mikecin Drazic, A.M., Mayakonda, A., Lutsik, P., Veith, C., et al. (2023). The GAPDH redox switch safeguards reductive capacity and enables survival of stressed tumour cells. Nat Metab 5. 10.1038/s42255-023-00781-3.

[58] Niimi N, Yako H, Takaku S, Chung SK, Sango K. Aldose Reductase and the Polyol Pathway in Schwann Cells: Old and New Problems. Int J Mol Sci. 2021 Jan 21;22(3):1031. doi: 10.3390/ijms22031031.

[59] Zhang, Z., Liew, C.W., Handy, D.E., Zhang, Y., Leopold, J.A., Hu, J., Guo, L., Kulkarni, R.N., Loscalzo, J., and Stanton, R.C. (2010). High glucose inhibits glucose-6-phosphate dehydrogenase, leading to increased oxidative stress and β-cell apoptosis. The FASEB Journal 24. 10.1096/fj.09-136572.

[60] Torrente, L., DeNicola, G.M. (2023). GAPDH redox redux—rewiring pentose phosphate flux. Nat Metab 5. 10.1038/s42255-021-00523-3

[61] DiBonaventura, M.D., Sadosky, A., Concialdi, K., Hopps, M., Kudel, I., Parsons, B., Cappelleri, J.C., Hlavacek, P., Alexander, A.H., Stacey, B.R., et al. (2017). The prevalence of probable neuropathic pain in the US: Results from a multimodal general-population health survey. J Pain Res 10. 10.2147/JPR.S127014.

[62] Segelcke D, Sondermann JR, Kappert C, Pradier B, Görlich D, Fobker M, Vollert J, Zahn PK, Schmidt M, Pogatzki-Zahn EM. Blood proteomics and multimodal risk profiling of human volunteers after incision injury: A translational study for advancing personalized pain management after surgery. Pharmacol Res. 2025 Feb;212:107580. doi: 10.1016/j.phrs.2025.107580.

