## Supplementary material for "Metabolic resilience rules sex-specific pain recovery during hormonal aging: a multi-omics analysis of neuropathy in mice": Table S1; Table S2; Table S3; Figure S1; Figure S2; Figure S3; Figure S4

^‡^Senior author

Lead contact

**Table S1. Assessment of the potential impact of the oestrus cycle on neuropathic pain response in 12-month-old mice.**

The fertility status of female mice was evaluated through vaginal smears prior to neuropathy induction to investigate its influence on mechanical threshold sensitivity (measured using an aesthesiometer, with values expressed in grams of force) and body weight (BW, expressed in grams). No significant differences were observed between females in the oestrus cycle and those not cycling, indicating that the fertility status did not affect these parameters.

| STRAIN & SEX | AGE | CYCLE | BW | AE |
| --- | --- | --- | --- | --- |
| CD1 F | 12M | CYCLE | 32,9 | 13,63 |
| CD1 F | 12M | CYCLE | 35,6 | 11,3 |
| CD1 F | 12M | CYCLE | 35,3 | 11,9 |
| CD1 F | 12M | CYCLE | 31 | 9,1 |
| CD1 F | 12M | CYCLE | 32,6 | 11,5 |
| CD1 F | 12M | NO CYCLE | 43,2 | 10,7 |
| CD1 F | 12M | NO CYCLE | 36,2 | 10,86 |
| CD1 F | 12M | NO CYCLE | 51,9 | 12,8 |
| CD1 F | 12M | NO CYCLE | 48,9 | 10,2 |
| CD1 F | 12M | CYCLE | 61,9 | 12,86 |

**Table S2.** **Whole blood** **AAs, C0, and ACCs of male and female mice in BL and D7 condition at 12 months of age (12M).** Data are expressed in μmol L^-1^ extracted sample. Differences between the study groups were assessed by 2-factor ANOVA. p-values lower than 0.050 are shown in bold.

|  |  | **BL Female, mean (SD)^a^,**  **μmol L^-1^** | | **D7 Female, mean (SD)^a^,**  **μmol L^-1^** | | **BL Male, mean (SD)^a^,**  **μmol L^-1^** | | **D7 Male, mean (SD)^a^,**  **μmol L^-1^** | | **Two way-ANOVA test, p^b^** | | |
| --- | --- | --- | --- | --- | --- | --- | --- | --- | --- | --- | --- | --- |
| **Metabolite** | | |  |  |  | |  | | **Gender** | | **Time (pre/post CCI)** | **Gender*Time (pre/post CCI)** |
| **C0** | | | 32 (7) | 33 (7) | 28 (8) | | 30 (7) | | 0,20 | | 0,46 | 0,91 |
| **C2** | | | 32 (9) | 33 (10) | 18 (3) | | 25 (8) | | **0,001** | | 0,13 | 0,31 |
| **C3** | | | 0.7 (0.3) | 1.0 (0.6) | 0.5 (0.2) | | 0.8 (0.3) | | 0,28 | | **0,002** | 0,49 |
| **C4** | | | 0.4 (0.1) | 0.4 (0.1) | 0.34 (0.08) | | 0.3 (0.1) | | **0,005** | | 0,93 | 0,28 |
| **C5** | | | 0.21 (0.08) | 0.3 (0.1) | 0.15 (0.07) | | 0.24 (0.08) | | 0,28 | | **0,003** | 0,17 |
| **C4OH/C3DC** | | | 0.20 (0.08) | 0.3 (0.2) | 0.20 (0.09) | | 0.2 (0.1) | | 0,34 | | 0,23 | 0,38 |
| **C6** | | | 0.09 (0.04) | 0.08 (0.03) | 0.08 (0.03) | | 0.08 (0.04) | | 0,72 | | 0,96 | 0,52 |
| **C5OH/C4DC** | | | 0.4 (0.1) | 0.5 (0.5) | 0.4 (0.1) | | 0.4 (0.3) | | **0,002** | | 0,79 | 0,10 |
| **C5DC/C6OH** | | | 0.3 (0.3) | 0.3 (0.1) | 0.17 (0.09) | | 0.3 (0.2) | | 0,55 | | 0,42 | 0,16 |
| **C8** | | | 0.09 (0.02) | 0.08 (0.04) | 0.07 (0.03) | | 0.09 (0.04) | | 0,9 | | 0,81 | 0,17 |
| **C6DC** | | | 0.013 (0.07) | 0.1 (0.1) | 0.2 (0.1) | | 0.2 (0.1) | | 0,17 | | 0,82 | 0,64 |
| **C10** | | | 0.06 (0.03) | 0.06 (0.02) | 0.05 (0.03) | | 0.08 (0.03) | | 0,58 | | 0,18 | 0,24 |
| **C12** | | | 0.08 (0.04) | 0.08 (0.04) | 0.08 (0.03) | | 0.10 (0.05) | | 0,38 | | 0,49 | 0,54 |
| **C14:1** | | | 0.10 (0.05) | 0.09 (0.03) | 0.10 (0.04) | | 0.10 (0.04) | | 0,53 | | 0,86 | 0,97 |
| **C14** | | | 0.3 (0.1) | 0.23 (0.08) | 0.3 (0.1) | | 0.3 (0.2) | | 0,10 | | 0,98 | 0,33 |
| **C14OH** | | | 0.04 (0.02) | 0.05 (0.03) | 0.06 (0.03) | | 0.06 (0.03) | | 0,17 | | 0,95 | 0,95 |
| **C16:1** | | | 0.19 (0.07) | 0.16 (0.05) | 0.21 (0.08) | | 0.20 (0.08) | | 0,086 | | 0,49 | 0,67 |
| **C16** | | | 1.3 (0.3) | 1.2 (0.2) | 1.2 (0.2) | | 1.5 (0.4) | | 0,41 | | 0,48 | 0,052 |
| **C16OH** | | | 0.11 (0.06) | 0.11 (0.06) | 0.14 (0.07) | | 0.16 (0.07) | | 0,064 | | 0,72 | 0,68 |
| **C18:2** | | | 0.21 (0.06) | 0.21 (0.04) | 0.18 (0.05) | | 0.24 (0.09) | | >0.99 | | 0,19 | 0,24 |
| **C18:1** | | | 0.6 (0.1) | 0.6 (0.1) | 0.5 (0.1) | | 0.7 (0.2) | | 0,98 | | 0,28 | 0,2 |
| **C18** | | | 0.4 (0.1) | 0.38 (0.09) | 0.35 (0.08) | | 0.4 (0.1) | | 0,92 | | 0,59 | 0,29 |
| **C18OH** | | | 0.07 (0.04) | 0.07 (0.03) | 0.10 (0.03) | | 0.10 (0.02) | | **0,016** | | 0,49 | 0,79 |

^a^ Standard deviation; ^b^ 95% confidence level; ^c^ 4 months of age; ^d^ 12 months of age.

*(continued)*

|  |  | **BL Female, mean (SD)^a^,**  **μmol L^-1^** | | **D7 Female, mean (SD)^a^,**  **μmol L^-1^** | | **BL Male, mean (SD)^a^,**  **μmol L^-1^** | | **D7 Male, mean (SD)^a^,**  **μmol L^-1^** | | **Two way-ANOVA test, p^b^** | | |
| --- | --- | --- | --- | --- | --- | --- | --- | --- | --- | --- | --- | --- |
| **Metabolite** | | |  |  |  | |  | | **Gender** | | **Time (pre/post CCI)** | **Gender*Time (pre/post CCI)** |
| **Short-chain ACCs** | | | 32 (9) | 33 (10) | 19 (3) | | 25 (8) | | **0,001** | | 0,13 | 0,32 |
| **Odd-chain ACCs** | | | 0.9 (0.3) | 1.2 (0.6) | 0.7 (0.2) | | 1.1 (0.4) | | 0,22 | | **<0.001** | 0,33 |
| **3-Hydroxy/Di-Carboxy ACCs** | | | 1.3 (0.4) | 1.4 (0.3) | 1.2 (0.3) | | 1.3 (0.3) | | 0,49 | | 0,38 | 0,9 |
| **Unsaturated chain ACCs** | | | 1.1 (0.3) | 1.1 (0.2) | 1.0 (0.3) | | 1.2 (0.4) | | 0,61 | | 0,55 | 0,33 |
| **Saturated chain ACCs** | | | 2.2 (0.6) | 2.0 (0.3) | 2.0 (0.5) | | 2.5 (0.7) | | 0,38 | | 0,55 | 0,10 |
| **Pro** | | | 48 (17) | 61 (26) | 54 (24) | | 62 (17) | | 0,60 | | 0,18 | 0,74 |
| **Val** | | | 67 (22) | 74 (21) | 65 (18) | | 73 (15) | | 0,75 | | 0,23 | 0,94 |
| **Leu/Ile/Pro-OH** | | | 166 (26) | 177 (51) | 151 (34) | | 198 (39) | | 0,86 | | **0,009** | 0,086 |
| **Orn** | | | 131 (78) | 186 (108) | 198 (129) | | 134 (60) | | 0,78 | | 0,89 | 0,092 |
| **Met** | | | 31 (10) | 36 (21) | 26 (10) | | 30 (13) | | 0,26 | | 0,32 | 0,91 |
| **Phe** | | | 38 (5) | 42 (8) | 34 (6) | | 42 (5) | | 0,37 | | **0,008** | 0,38 |
| **Arg** | | | 65 (19) | 63 (20) | 51 (13) | | 58 (20) | | 0,074 | | 0,73 | 0,46 |
| **Cit** | | | 65 (22) | 72 (24) | 58 (27) | | 45 (19) | | **0,022** | | 0,73 | 0,20 |
| **Tyr** | | | 74 (14) | 78 (16) | 68 (13) | | 88 (23) | | 0,77 | | 0,072 | 0,21 |
| **Gly** | | | 723 (419) | 917 (460) | 1203 (554) | | 857 (468) | | 0,097 | | 0,674 | 0,144 |
| **Ala** | | | 124 (90) | 122 (97) | 146 (94) | | 122 (76) | | 0,65 | | 0,70 | 0,75 |
| **Ser** | | | 4 (2) | 4 (3) | 4 (3) | | 5 (3) | | 0,50 | | 0,44 | 0,96 |
| **Thr** | | | 18 (9) | 17 (7) | 15 (6) | | 19 (9) | | >0.99 | | 0,54 | 0,29 |
| **Asn** | | | 3 (1) | 3.4 (0.8) | 3 (2) | | 4 (2) | | 0,23 | | 0,084 | 0,54 |
| **Asp** | | | 6 (2) | 6 (2) | 6 (1) | | 7 (2) | | 0,90 | | 0,090 | 0,15 |
| **Lys/Gln** | | | 10444 (4606) | 12599 (5540) | 11297 (7061) | | 6417 (1617) | | 0,073 | | 0,46 | 0,068 |
| **Glu** | | | 23 (3) | 21 (7) | 23 (10) | | 29 (13) | | 0,074 | | 0,52 | 0,26 |
| **His** | | | 302 (202) | 311 (142) | 224 (222) | | 288 (159) | | 0,47 | | 0,43 | 0,56 |

^a^ Standard deviation; ^b^ 95% confidence level; ^c^ 4 months of age; ^d^ 12 months of age.

**Table S3.** **Multiple post-hoc comparisons between whole blood AAs, C0, and ACCs of male and female mice in BL and D7 condition at 12 months of age (12M).** The data are p values obtained by Fischer test. p-values lower than 0.050 are shown in bold.

|  | **Fisher post hoc test, p^a^** | | | |
| --- | --- | --- | --- | --- |
| **Metabolite** | **BL Male *vs* D7 Male** | **BL Male *vs* BL Female** | **D7 Male *vs* D7 Female** | **BL Female *vs* D7 Female** |
| **C0** | 0,54 | 0,34 | 0,38 | 0,65 |
| **C2** | 0,082 | **0,004** | **0,048** | 0,71 |
| **C3** | **0,008** | 0,30 | 0,61 | 0,061 |
| **C4** | 0,41 | 0,23 | **0,004** | 0,48 |
| **C5** | **0,003** | 0,17 | 0,89 | 0,18 |
| **C4OH/C3DC** | 0,81 | 0,87 | 0,14 | 0,15 |
| **C6** | 0,68 | 0,53 | 0,91 | 0,63 |
| **C5OH/C4DC** | 0,17 | 0,22 | **0,002** | 0,31 |
| **C5DC/C6OH** | 0,13 | 0,11 | 0,42 | 0,66 |
| **C8** | 0,25 | 0,44 | 0,35 | 0,41 |
| **C6DC** | 0,62 | 0,48 | 0,21 | 0,86 |
| **C10** | 0,080 | 0,43 | 0,13 | 0,89 |
| **C12** | 0,36 | 0,86 | 0,28 | 0,96 |
| **C14:1** | 0,88 | 0,63 | 0,68 | 0,92 |
| **C14** | 0,50 | 0,84 | **0,038** | 0,48 |
| **C14OH** | >0.99 | 0,31 | 0,35 | 0,93 |
| **C16:1** | 0,85 | 0,41 | 0,10 | 0,44 |
| **C16** | 0,063 | 0,24 | **0,027** | 0,35 |
| **C16OH** | 0,59 | 0,27 | 0,12 | 0,97 |
| **C18:2** | 0,083 | 0,40 | 0,40 | 0,92 |
| **C18:1** | 0,099 | 0,22 | 0,23 | 0,88 |
| **C18** | 0,26 | 0,40 | 0,48 | 0,70 |
| **C18OH** | 0,76 | 0,11 | 0,056 | 0,49 |
| **Short-chain ACCs** | 0,083 | **0,004** | 0,045 | 0,70 |
| **Odd-chain ACCs** | **0,002** | 0,21 | 0,62 | **0,039** |
| **3-Hydroxy/Di-Carboxy ACCs** | 0,47 | 0,55 | 0,71 | 0,59 |
| **Unsaturated chain ACCs** | 0,27 | 0,55 | 0,20 | 0,79 |
| **Saturated chain ACCs** | 0,12 | 0,38 | **0,043** | 0,44 |

^a^ 95% confidence level.

*(continued)*

|  | **Fisher post hoc test, p^a^** | | | |
| --- | --- | --- | --- | --- |
| **Metabolite** | **BL Male *vs* D7 Male** | **BL Male *vs* BL Female** | **D7 Male *vs* D7 Female** | **BL Female *vs* D7 Female** |
| **Pro** | 0,46 | 0,53 | 0,91 | 0,24 |
| **Val** | 0,37 | 0,78 | 0,87 | 0,42 |
| **Leu/Ile/Pro-OH** | **0,004** | 0,46 | 0,32 | 0,44 |
| **Orn** | 0,19 | 0,11 | 0,21 | 0,26 |
| **Met** | 0,52 | 0,46 | 0,38 | 0,43 |
| **Phe** | **0,013** | 0,20 | 0,98 | 0,16 |
| **Arg** | 0,44 | 0,062 | 0,50 | 0,78 |
| **Cit** | 0,25 | 0,48 | **0,011** | 0,49 |
| **Tyr** | **0,036** | 0,31 | 0,16 | 0,67 |
| **Gly** | 0,18 | **0,011** | 0,73 | 0,45 |
| **Ala** | 0,61 | 0,53 | 0,99 | 0,96 |
| **Ser** | 0,56 | 0,65 | 0,62 | 0,60 |
| **Thr** | 0,24 | 0,43 | 0,43 | 0,75 |
| **Asn** | 0,10 | 0,60 | 0,24 | 0,41 |
| **Asp** | **0,032** | 0,57 | 0,46 | 0,84 |
| **Lys/Gln** | 0,072 | 0,67 | **0,006** | 0,41 |
| **Glu** | 0,21 | 0,80 | **0,026** | 0,72 |
| **His** | 0,33 | 0,44 | 0,81 | 0,88 |

^a^ 95% confidence level.

**Table S4. DIFFERENTIAL PROTEOMIC ANALYSIS DETAILS.** A total of 406 DEPs were selected from four groups of mice (females in baseline condition – F_BL; females at day 7 after CCI – F_D7; males in baseline condition – M_BL; males at day 7 after CCI – M_D7), with a maximum fold change (MFC) in protein expression levels set at MFC ≥ 1.5 and statistically significant observations (ANOVA p-value ≤ 0.05).
Details of the analysis are provided in the attached Excel file named “Table 4 Differential Proteins”, due to the large size of the dataset, the presence of multiple Excel sheets, and the extensive list of proteins**.**

**Figure S1**. Volcano Scatter Plots of pair comparison. Significance is plotted on the y axes ( -Log10 of p-value ) and  fold-change (Log2 of Fold Change) plotted on the x axes. Left side of the Volcano represent proteins downregulated and while the right side indicate proteins upregulated. VolcaNoseR is a web app for creating, exploring, labeling and sharing volcano plots (26) https://huygens.science.uva.nl/VolcaNoseR/

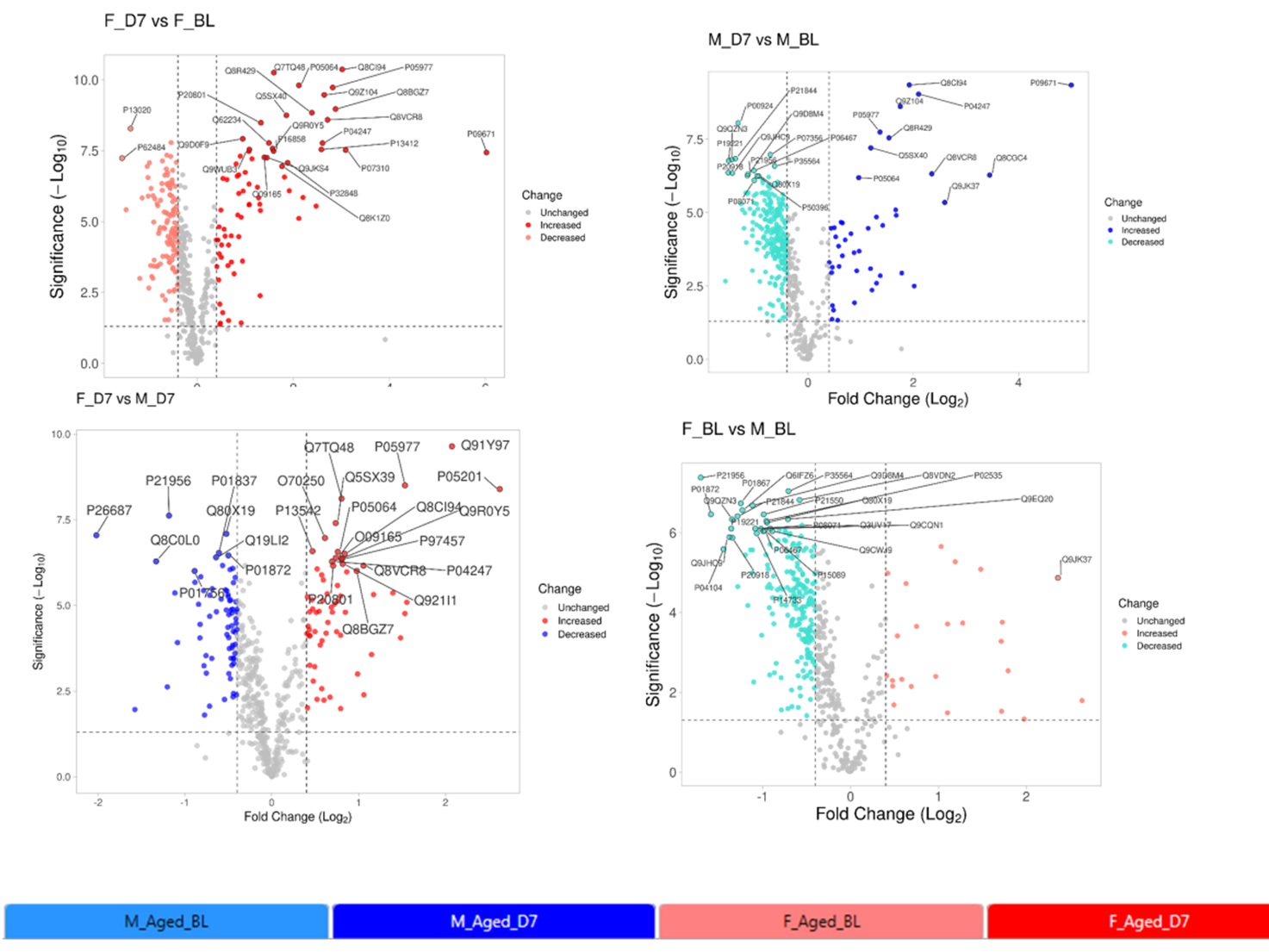

Figure S2. Most Relevant Pathway enriched by the DEPs in females baseline condition vs males baseline condition (FBLvsMBL) sorted by p-value

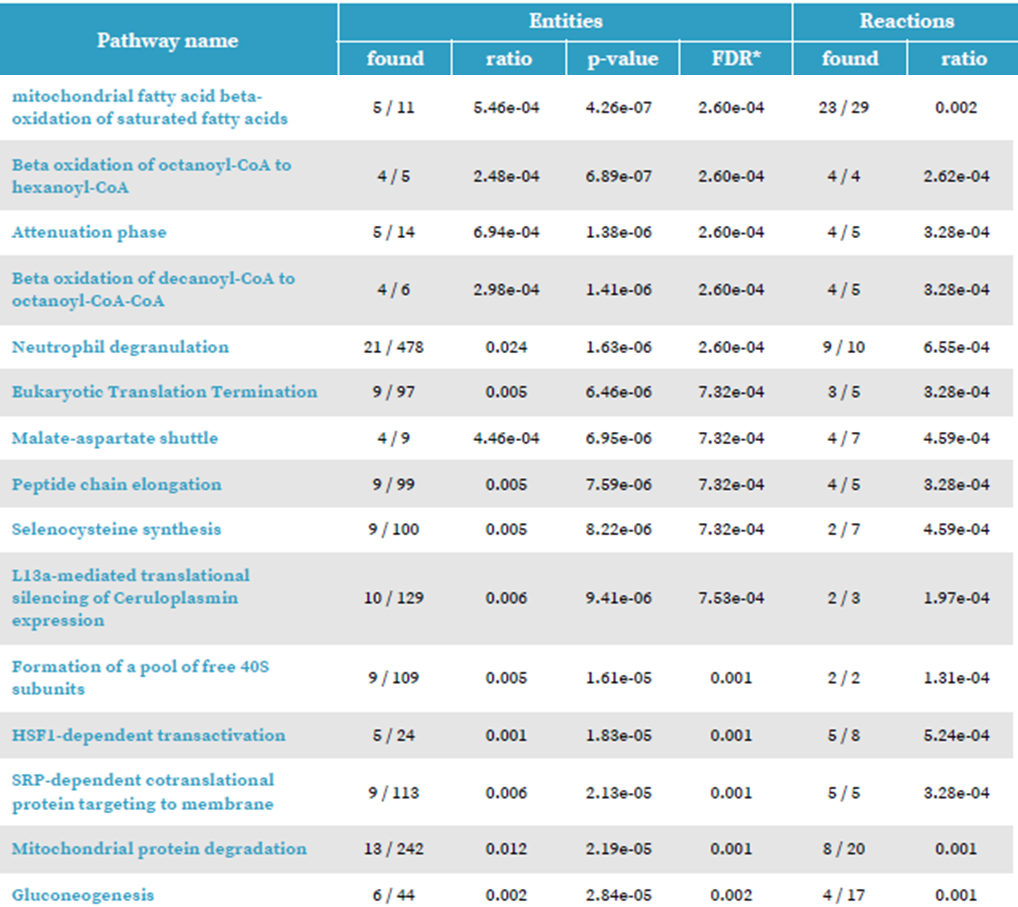

Figure S3. Most Relevant Pathway enriched by the DEPs in females 7 days after CCI vs males 7 days after CCI (FD7vsMD7) sorted by p-value

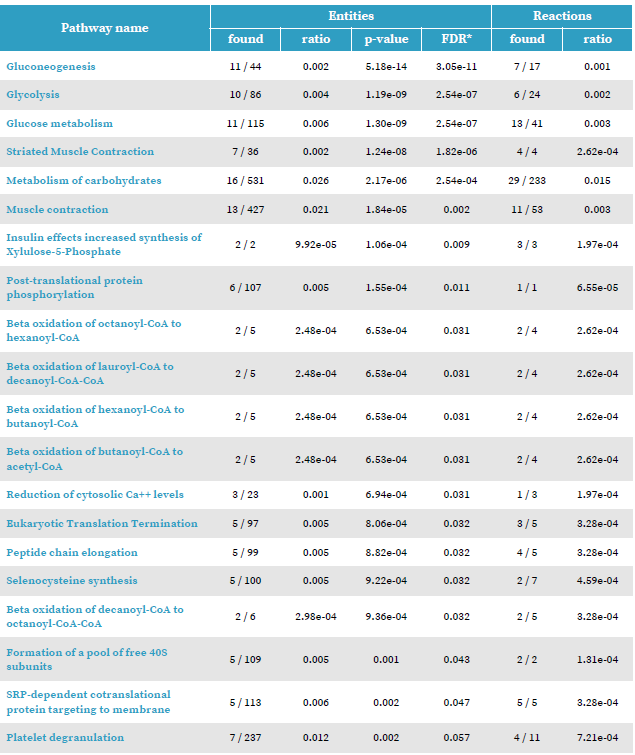

Figure S4. Leptin serum levels (ELISA) measured in naïve animals and male and female mice 7 days after CCI.

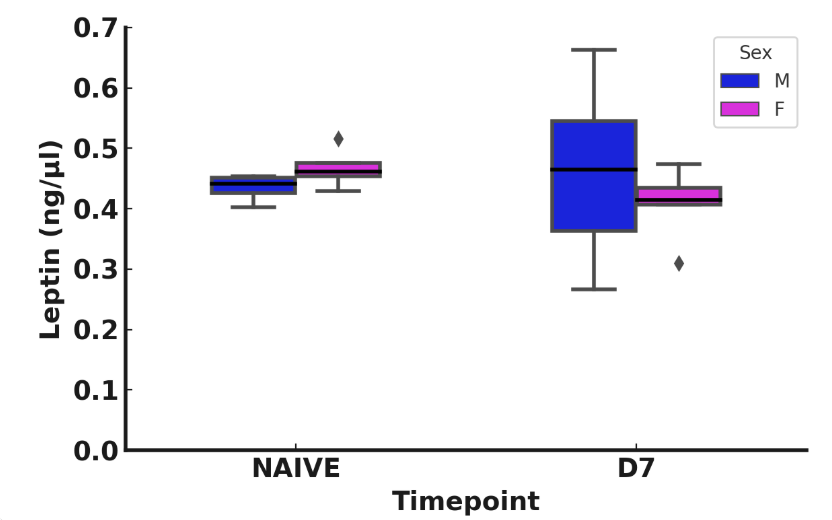
